# Mechanistic Language Modeling and Oxygenated 3D Screening Reveal Berberine and Enzalutamide Synergy in Resistant Prostate Cancer

**DOI:** 10.64898/2026.01.24.701539

**Authors:** Chih-Hui Lo, Katie Shi, Lina Kafadarian, Alexandra Bermudez, Johnny Diaz, Liam Edwards, Yunqi Hong, Ziyi Chen, Hyeonji Hwang, Weihong Yan, Alan Levinson, Robert Damoiseaux, Cho-Jui Hsieh, Tanya Stoyanova, Andrew S. Goldstein, Neil Y.C. Lin

**Affiliations:** Bioengineering Department, University of California Los Angeles, Los Angeles, CA, USA; College of Pharmacy, National Defense Medical University, Taipei, Taiwan; Mechanical and Aerospace Engineering Department, University of California Los Angeles, Los Angeles, CA, USA; Department of Molecular, Cell, and Developmental Biology, University of California Los Angeles, Los Angeles, CA, USA; Computer Science Department, University of California Los Angeles, Los Angeles, CA, USA; Department of Molecular and Medical Pharmacology, University of California Los Angeles, Los Angeles, CA, USA; Department of Chemistry and Biochemistry, University of California, Los Angeles, CA, USA; California NanoSystems Institute, University of California Los Angeles, Los Angeles, CA, USA; Eli and Edythe Broad Center of Regenerative Medicine and Stem Cell Research, University of California Los Angeles, Los Angeles, CA, USA; Jonsson Comprehensive Cancer Center, University of California Los Angeles, Los Angeles, CA, USA; Department of Urology, University of California Los Angeles, Los Angeles, CA, USA; Institute for Quantitative and Computational Biosciences, University of California Los Angeles, Los Angeles, CA, USA

**Author notes:** Corresponding Authors: Neil Y.C. Lin. **Conflict of Interest:** The authors declare no potential conflicts of interest.

## Abstract

Resistance to androgen receptor inhibitors remains a primary challenge in prostate cancer treatment, yet identifying synergistic co-therapies is hindered by immense combinatorial search spaces and the limited interpretability of predictive computation models. Here, we developed an integrated discovery-validation axis coupling knowledge-augmented large language models with oxygen-supplemented 3D spheroid assays. By leveraging inherent model stochasticity, our framework measures the degree of consensus across independent predictions to establish a formal metric for predictive accuracy. This principle enables high-throughput assessment of complex signaling crosstalk, yielding mechanistic rationales for all predictions and defining a high-confidence zone that minimizes experimental attrition. Utilizing this approach to screen 3,592 natural products, we identified a previously unrecognized synergy between berberine and enzalutamide that re-sensitizes resistant cells. Validation confirms that berberine perturbs the PI3K/AKT/mTOR and AMPK axes, a finding consistent with the mechanistic rationales computationally derived by the framework. Integrating interpretable AI with physiologically relevant 3D screening provides a scalable methodology for the rational discovery of synergistic therapies.

**Significance:** Integrating mechanistic AI with oxygenated 3D screening, we identify a novel berberine-enzalutamide synergy. This framework resolves complex signaling dependencies, providing a scalable, transparent methodology for the rational discovery of effective combination therapies.

## Introduction

Prostate cancer remains a primary driver of male cancer mortality globally^1^. While androgen deprivation therapy serves as the initial standard of care, tumors invariably progress to a castration-resistant state^2^. In this setting, next-generation androgen receptor (AR) signaling inhibitors, such as enzalutamide, provide survival benefits; however, clinical efficacy is frequently short-lived due to the rapid emergence of acquired resistance^3–5^. This resistance is driven by a highly plastic adaptive landscape where cancer cells engage compensatory signaling bypasses to maintain survival despite AR blockade^6–12^. As such, identifying synergistic drug combinations capable of co-inhibiting these bypass mechanisms is essential for improving patient outcomes^11,13,14^. Unfortunately, the high-dimensional nature of the combinatorial landscape makes exhaustive experimental screening intractable, even with high-throughput automation.

To address this hurdle, *in silico* screening has been adopted to facilitate the experimental pipeline, serving as a predictive filter to prioritize therapeutic candidates for downstream validation. Established deep learning architectures, including DeepSynergy^15^ and DeepDDs^16^, have demonstrated significant predictive utility by identifying patterns within structured chemical and biological datasets^17–21^. While these models provide high accuracy in statistical feature attribution^22,23^, their continued refinement increasingly depends on integrating mechanistic context to strengthen biological validation. This capability has been further expanded by recent Large Language Model (LLM) frameworks, such as CancerGPT^24^ and SynerGPT^25^, which leverage vast repositories of unstructured biomedical knowledge to bridge data gaps^26^. However, an ongoing challenge remains in distinguishing linguistic probability from biological truth. Without explicit uncertainty quantification^27–30^, computational leads may remain susceptible to high-probability hallucinations. Addressing these gaps is essential for ensuring that computational predictions can fully resolve the complex signaling dependencies of the resistant landscape.

While computational frameworks facilitate candidate prioritization, the experimental substrate remains the ultimate arbiter of translational utility. Two-dimensional monolayers offer high scalability, yet they frequently fail to recapitulate the complex spatial architecture and cell-cell interactions that drive drug resistance in vivo. This discrepancy necessitates the transition to three-dimensional (3D) models for more predictive screening^31^. Despite their advantages, the reliability of conventional 3D spheroids is often compromised by physical constraints. To accurately mimic physiological gradients and drug penetration barriers, spheroids must typically exceed 400–500 µm in diameter. However, at this scale, the model encounters a critical threshold: diffusion-limited oxygen levels begin to drive the formation of stagnant necrotic cores, potentially confounding experimental outcomes^32–36^. These cores are characterized by non-specific cell death and disrupted cell-cell junctions that diverge from drug-induced responses^35–38^. Such phenotypic confounders mask the specific signaling dependencies identified as synergistic targets. Eliminating these non-specific artifacts is therefore critical to ensure that drug-response measurements reflect true therapeutic sensitivity rather than model-induced stress^36,38^. Altogether, identifying viable therapeutic candidates hinges on the integration of high-viability experimental platforms with mechanistic screening models that offer transparent, predictive insights.

Here, we present a unified discovery-validation axis centered on MAESTRO (Mechanistic Augmentation and Ensemble Selection for Therapeutic Ranking Optimization). MAESTRO is a knowledge-augmented LLM framework that harnesses inherent model stochasticity to quantify predictive confidence and evaluate accuracy. This approach enables the complex analysis of signaling crosstalk at scale, generating quantifiable synergy predictions alongside evidence-based rationales. This principle establishes a high-confidence zone for candidate prioritization, mitigating downstream experimental attrition. To ensure the physiological relevance of these predictions, we coupled MAESTRO with an oxygen-supplemented 3D spheroid platform, eliminating the confounding effects of hypoxic signaling inherent to standard models. Using this integrated pipeline, we identified a previously unrecognized synergy between the natural product berberine and enzalutamide. While berberine has been noted for general anti-proliferative effects^39–41^, our platform resolved a specific, potent synergistic interaction driven by the concurrent modulation of the PI3K/AKT/mTOR and AMPK signaling axes. Molecular and transcriptomic analyses confirm that this dual-axis modulation effectively re-sensitizes resistant cells to AR blockade, corroborating the mechanistic rationales computationally hypothesized by the framework. Together, our integrated approach provides a platform for the discovery of combination therapies that may be difficult to resolve using conventional screening modalities.

## Materials and Methods

### Architecture of the MAESTRO framework

The MAESTRO pipeline constructs context-augmented prompts by integrating drug-MoA labels with disease-specific mechanistic contexts. For drug-MoA generation, we utilize PubTator3^42^ to retrieve biomedical articles, executing a Boolean search for each compound using the keywords: "mechanism", "target", "gene", "protein", "pathway" "signaling", "biological" "disease" "cancer", "activate", or "inhibit". These articles are then processed by OpenAI GPT-4o-mini (temperature = 1.0) to synthesize concise profiles of molecular targets, downstream signaling, and potential AR signaling crosstalk. Simultaneously, the disease-specific context is derived via a two-stage RAG workflow from 13 curated review articles on prostate cancer resistance. A structured prompt explicitly queries 12 conceptual categories including epigenetic modifiers, upstream/downstream regulators, and cellular phenotypes to ensure a comprehensive mechanistic framework that connects upstream regulators to downstream phenotypes.

To accommodate context window limitations and mitigate the performance degradation associated with excessive token loads, candidates were processed in batches of 50 compounds per prompt. We employ a balanced random sampling algorithm to ensure each candidate is evaluated with equivalent frequency against randomized cohorts, effectively mitigating batch-specific bias. Ranking is performed using OpenAI GPT-4o or Google Gemini 2.5 Flash (temperature = 1.0, max token: 2000, all other settings default). To quantify aleatoric uncertainty and stabilize the output, we implement an ensemble consensus strategy where each candidate undergoes a minimum of 100 independent prioritizations (with up to 400 runs for specific analyses). Final consensus rankings and uncertainty metrics are derived from the distribution of these iterations.

The workflow is formally defined as follows (Fig. 1a): let *D* = {*d*_1_*, d*_2_*, . . . , d_n_*} denote the library of candidate drugs. In each independent sampling iteration *t*, a subset *S_t_* ⊂ *D* is constructed via the balanced random sampling strategy. The synergy assessment is modeled as a ranking function *R_t_* : *S_t_* → {1*,…,* 50}, which represents the mechanistic reasoning process mapping a drug’s context to a specific rank. Consequently, *R_t_*(*i*) denotes the rank assigned to drug *d_i_* in sampling run *t*. The final consensus prioritization is quantified by the mean rank 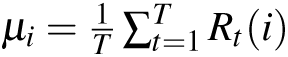 , where *T* is the total number of iterations. The aleatoric uncertainty is captured by the standard deviation 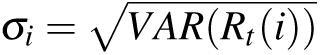.

**Figure 1.**
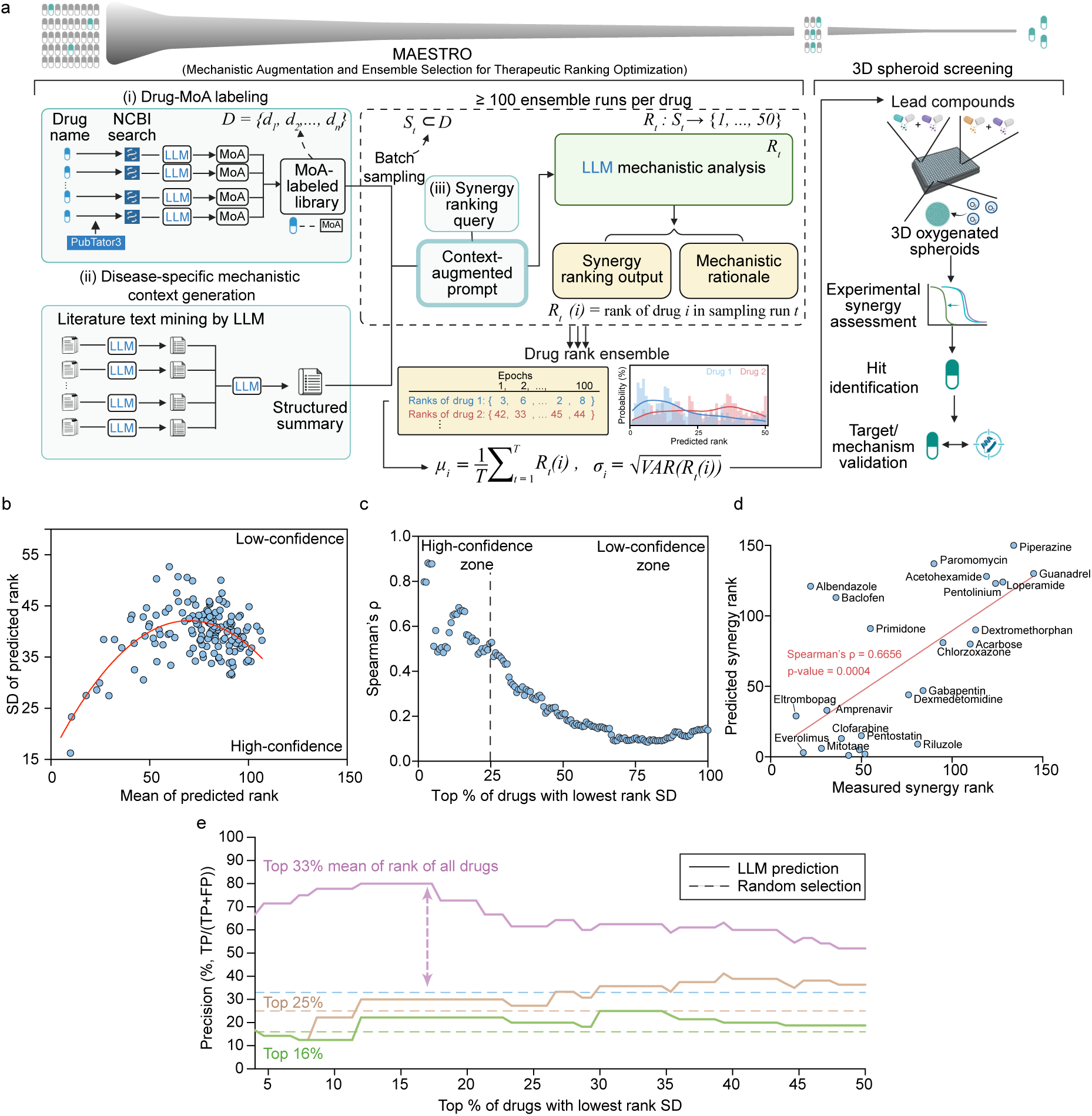
MAESTRO leverages mechanistic augmentation and ensemble consensus for high-confidence drug synergy predictions. **(a)** To identify lead compound predictions, the MAESTRO pipeline begins by generating (i) LLM-derived compound mechanism of action (MoA) labels grounded in NCBI research articles. It also generates (ii) disease-specific contexts mined from the literature. Both the Drug MoA labeled library and the disease-specific mechanistic context are used as input to another LLM. Utilizing randomized batch sampling, subsets of compound labels are iteratively synthesized with (iii) a synergy-ranking query to construct context-augmented prompts. These prompts are processed by LLMs to accumulate a minimum of 100 ensemble runs per compound, where the mean rank summarizes consensus prioritization and rank standard deviation (SD) quantifies aleatoric uncertainty. High-confidence lead candidates are finally validated in an oxygen-supplemented 3D prostate cancer spheroid model. *D*, whole compound library; *S_t_* , randomly sampled subset at iteration *t*; *R_t_* (*i*), rank of compound *i* at iteration *t*. **(b)** Scatter plot illustrating the inverted parabolic relationship between MAESTRO-predicted mean rank and rank standard deviation (SD) for 150 AR inhibitor-anchored drug pairs from the DrugComb database (400 runs per compound). Low rank SD signifies high-confidence predictions, whereas high rank SD denotes low confidence. The red line represents a second-order polynomial fit. **(c)** Spearman’s rank correlation (*ρ*) between predicted and experimental rankings versus the percentage of drugs included, sorted by ascending rank SD. **(d)** Predicted synergy ranking versus Predicted synergy rank for the 24 representative drugs exhibiting the lowest rank SD (high-confidence). The red line represents the best fit. **(e)** Screening precision profiles plotted against the cumulative percentage of drug pairs included, ordered by ascending rank SD. Solid lines represent precision for varying candidate selection thresholds (top 16%–33% based on mean rank), while horizontal dashed lines indicate the corresponding random selection baselines. TP, true positive; FP, false positive.

### MAESTRO’s prediction characterization

In our verification workflow, we cross-referenced the 95 prioritized PKIs against published records to distinguish pre-existing knowledge from novel inferences. Two independent researchers with biomedical domain expertise systematically searched Google Scholar for each candidate using the query string: “[Candidate Name] AND Enzalutamide”. We restricted the manual screening scope to the top 100 results sorted by relevance. Validated synergy reports were defined based on the following inclusion criteria: (i) both drug names were co-present in the text; and (ii) the article explicitly reported a synergistic interaction or combination benefit in either *in vitro* or *in vivo* settings.

To evaluate the relationship between target engagement directness and MAESTRO’s prioritization, we systematically processed 127 previously classified PI3K/AKT/mTOR inhibitors using the LLM alongside their MoA labels. We instructed the model to assign a “Target Involvement Score” (scale 1–10) based on the specific definitions provided in the prompt. The exact scoring criteria provided to the LLM were as follows: "PI3K–AKT–mTOR Target Involvement Score (1 to 10). This score measures how strongly a drug directly acts on the PI3K–AKT–mTOR signaling axis, based only on its known mechanism of action, target descriptions, and how it is classified in the literature. How to assign the score: Score 9–10: The drug is a core, high-confidence inhibitor of PI3K, AKT, or mTOR. It is clearly described as an AKT inhibitor, PI3K inhibitor, mTOR inhibitor, or dual PI3K/mTOR inhibitor. These drugs are widely recognized and categorized in reviews as canonical PI3K-axis inhibitors. Score 7–8: The drug strongly inhibits the PI3K pathway but not exclusively. It may target PI3K-axis kinases along with other kinases, or inhibit a very close upstream regulator. PI3K/AKT/mTOR inhibition is still a major part of its pharmacology but shares space with other targets. Score 5–6: The drug is not labeled as a PI3K/AKT/mTOR inhibitor, but multiple studies show that it meaningfully suppresses this pathway under some conditions. These effects tend to be indirect or context-dependent. Score 3–4: The drug has weak or occasional effects on the PI3K axis. It may reduce AKT phosphorylation only at high concentrations or in specific models, and this is not a major feature of its mechanism. Score 1–2: The drug has no meaningful involvement with the PI3K–AKT–mTOR axis. Its primary targets belong to unrelated biology (e.g., metabolic enzymes, immune receptors, malaria proteins, transcription modulators, Rac1/ROCK, GCN2, etc.)."

To perform a semantic analysis of the mechanistic reasoning, we first encoded the LLM-generated mechanistic justifications (*n* = 100 per compound) into high-dimensional dense vectors using the all-MiniLM-L6-v2 SentenceTransformer model^43^. We then projected these semantic embeddings into a two-dimensional space to visualize latent clustering patterns using t-SNE (perplexity: 30; initialization: PCA) and PCA. All visualizations were generated with a fixed random state of 40. For word cloud visualization, we stratified the 127 PI3K/AKT/mTOR inhibitors into three cohorts: Top (Rank 1–30), Middle (Rank 50–80), and Bottom (Rank 100–127). We aggregated the mechanistic reasoning texts (*n* = 100 per compound) within each cohort and removed standard English stopwords and domain-specific generic terms (e.g., “inhibitor”, “kinase”, “signaling”). The WordCloud Python library was used to generate visualizations, scaling term size proportionally to frequency within the aggregated corpus.

To develop our LLM-driven evidence scoring for MoA annotations, we established a 0–5 ordinal Evidence Score to correlate the degree of mechanistic evidence in drug-MoA labels with MAESTRO rankings. To minimize self-validation bias, we employed Google Gemini 2.5 Flash as an independent evaluator, distinct from the annotating GPT-4o, to analyze the natural product (NP) drug-MoA labels. The model assigned scores across 12 key signaling pathways and molecular targets, yielding a 12-dimensional feature vector based on the following strict criteria: "Score 0: No evidence. Criteria: No evidence that indicates any interaction; Score 1: Speculative (In Silico). Criteria: Computational prediction, molecular docking, or hypothesis only. No wet-lab data; Score 2: Indirect (Associative). Criteria: Secondary effects (e.g., mRNA expression changes), correlations, or general stress responses without a clear mechanism; Score 3: Functional (In Vitro). Criteria: Demonstrated change in cellular signaling (e.g., Western blot showing reduced phosphorylation or nuclear translocation) in cell culture; Score 4: Direct (Biochemical). Criteria: Proof of physical binding or direct enzymatic inhibition (e.g., kinase assays, ligand displacement); Score 5: Validated (In Vivo/Clinical). Criteria: Confirmed mechanism of action driving efficacy in animal models (xenografts) or human clinical trials."

To conduct the multivariate regression and SHAP interpretability analysis, we grouped the NP library into four subsets (top 100, top 500, top 1,000, and complete library) based on the lowest SD of MAESTRO-predicted ranks. Independent multivariate regression analyses were performed for each subset using the Gradient Boosted Trees (GBT) algorithm^44^ implemented in scikit-learn. The GBT model utilized an ensemble of 300 decision trees with a learning rate of 0.05 and a maximum tree depth of 3 to balance complexity and generalization. A fixed random state was applied for reproducibility. We evaluated model performance by calculating the Coefficient of Determination (*R*^2^) and Root Mean Square Error (RMSE) between MAESTRO-predicted and GBT-predicted ranks. Feature contributions were quantified using SHapley Additive exPlanations (SHAP)^45^. We computed local SHAP values using the TreeExplainer method (shap Python package) and determined global feature importance by calculating the mean absolute SHAP value for each evidence category.

### Cell culture and generation of oxygenated PCa spheroids

LNCaP and 16D^CRPC^ cells were cultured in RPMI 1640 media (Gibco, Cat# 11875093) with 10% fetal bovine serum (FBS; Gibco, Cat# 26140079) and 1% penicillin/streptomycin (P/S; Gibco, Cat# 15140122). 22RV1 cells were cultured in RPMI 1640 ATCC modification (Gibco, Cat# A1049101) with 10% FBS and 1% P/S. VCaP cells were cultured in DMEM (Gibco, Cat# 11995040) with 10% FBS and 1% P/S. C4-2B MDVR cells^46^ were received from Dr. Allen Gao, UC Davis, and cultured in RPMI 1640 media with 10% FBS, 1% P/S, and 20 *µ*M enzalutamide (Apexbio, Cat# A3003). All 2D cells and 3D spheroids were maintained at 37^◦^C, 5% CO_2_. Media changes were performed every 2–3 days, with subculturing conducted when cells reached to 80–90% confluence. All cell lines tested negative for mycoplasma contamination by PCR and were used for experiments at low passage numbers.

To generate PCa spheroids with sizes of 500-600 *µ*m, 22RV1, C4-2B MDVR, LNCaP, 16D^CRPC^, or VCaP cells were seeded at 10,000, 1,000, 2,000, 3,000, and 15,000 cells per well, respectively, into 96-well cell-repellent V-bottom plates (Greiner Bio-One, Cat# 651970). Immediately after seeding, plates were centrifuged at 100 × g for 1 min to promote aggregation. Aggregates were cultured at 21% O_2_ for 48 h to form compact spheroids. Each spheroid per well was then gently transferred to 96- or 384-well ultra-low-attachment round-bottom plates (Corning, Cat# 4515 and 4516, respectively) for optimal imaging quality. For oxygen-level optimization assays, spheroids were cultured under 21%, 40%, 60%, and 95% O_2_ for 72 h prior to downstream characterization using sealed sub-chambers purged with pre-mixed gas cylinders. Following the identification of 40% O_2_ as the optimal condition, a tri-gas incubator (Thermo Scientific) was employed to maintain 40%O_2_ for all subsequent drug-screening assays. In drug-screening assays, PCa spheroids were treated with compounds and maintained at 40% O_2_ for 72 h.

### Immunofluorescence staining

Following 72 h incubation at their respective oxygen levels, PCa spheroids were fixed in 4% paraformaldehyde (Thermo Scientific Chemicals, Cat# 043368-9M), embedded in OCT compound, and snap-frozen. Cryosections (12 *µ*m) were prepared, then permeabilized and blocked in PBS containing 2% donkey serum and 0.25% Triton X-100 (Sigma-Aldrich, Cat# X100) at room temperature for 30 min. After PBS washes, primary antibodies against Ki-67 (Cell Signaling Technology, RRID: AB_2797703), cleaved caspase-3 (CC3; Cell Signaling Technology, RRID: AB_2341188), E-cadherin (Proteintech, RRID: AB_10697811), and *γ*-H2AX (Thermo Fisher Scientific, RRID: AB_2573048) were applied in PBS with 0.125% Triton X-100 and 0.5% bovine serum albumin (BSA; Fisher BioReagents, Cat# BP671-10) and incubated at 4^◦^C overnight. Primary antibodies were diluted according to the manufacturers’ instructions. Following PBS washes, fluorophore-conjugated secondary antibodies (1:1000) and 10 *µ*g mL^−1^ Hoechst 33342 (Hoechst; Invitrogen Cat# H1399) were applied and incubated at room temperature for 1 h in the dark. Slides were mounted with ProLong Diamond Antifade Mountant (Invitrogen, Cat# 36970) and imaged on a Revolution fluorescence microscope (ECHO) using identical acquisition settings across groups. For quantitative analysis, image data was processed using QuPath software^47^. Cells were detected from the Hoechst channel using the Cell Detection tool (Gaussian sigma, 2.0 *µ*m; minimum/maximum object area, 10 – 40,000 *µ*m^2^; cell expansion, 5.0 *µ*m to define whole-cell ROIs). Biomarker positivity was assigned using the Object Classifier based on nuclear mean intensity for Ki-67 and *γ*-H2AX, and expanded whole-cell mean intensity for CC3. Positivity was reported as the ratio of positive samples to the total number analyzed, with identical detection parameters and thresholds applied across all groups.

For 2D monolayer cultures, 22RV1 cells were seeded into 12-well removable chamber slides (ibidi, Cat# 81201) at a density of 140,000 cells cm^−2^ and cultured for 24 h at 37^◦^C with 5% CO_2_. Cells were treated with vehicle, 20 *µ*M enzalutamide (Enza), 5 *µ*M berberine (BBR; MedChemExpress, Cat# HY-N0716B), or the combination (Combo) for 8 h. Following treatment, cells were fixed in 4% paraformaldehyde containing phosphatase inhibitor cocktail (Cell Signaling Technology, Cat# 5870S) for 30 min on ice, then permeabilized and blocked in PBS containing 2% donkey serum and 0.25% Triton X-100 at room temperature for 30 min. After PBS washes, primary antibodies against phospho-S6 (p-S6; Cell Signaling Technology, RRID: AB_10694233), total S6 (Cell Signaling Technology, RRID :AB_331355), phospho-AMPK*α* (p-AMPK*α*; Cell Signaling Technology, RRID :AB_2799368), and total AMPK*α* (Cell Signaling Technology, RRID :AB_10622186) were applied in PBS with 0.125% Triton X-100 and 0.5% BSA and incubated at 4^◦^C overnight. Primary antibodies were diluted according to the manufacturer’s instructions. Following PBS washes, fluorophore-conjugated secondary antibodies (1:1000) and 10 *µ*g mL^−1^ Hoechst 33342 were applied and incubated at room temperature for 1 h in the dark. Slides were mounted with ProLong Diamond Antifade Mountant and imaged on a Zeiss Axio Observer Z1 (Carl Zeiss) inverted microscope equipped with an NL5+ line-scanning confocal module (Confocal NL). Image acquisition was performed using Micro-Manager software with identical settings across all groups. For quantitative analysis, image data were processed using a workflow integrating Cellpose and ImageJ. Nuclei were segmented from the Hoechst channel using Cellpose (model type, ‘nuclei’) to obtain cell counts per image. Biomarker expression was quantified in ImageJ by generating above-threshold masks for each channel. The integrated density was computed within the mask (mean intensity × mask area) and normalized by the nuclear count to yield mean fluorescence intensity per cell (MFI/cell). Identical segmentation parameters and intensity thresholds were applied across all treatment conditions.

### Live/dead staining

Following 72 h incubation at their respective oxygen levels, PCa spheroids were washed twice with serum-free medium and incubated with 2 *µ*M Calcein-AM (Biotium; live-cell stain, Cat# 80011-2), 5 *µ*M propidium iodide (PI; Biotium; dead-cell stain, Cat# 40017), and 10 *µ*g mL^−1^ Hoechst 33342 at 37^◦^C for 3 h. Spheroids were then imaged on a Revolution fluorescence microscope (ECHO) using identical acquisition settings across groups. For quantitative analysis, after setting a threshold to subtract the background, Calcein-AM and propidium iodide total intensity of the spheroids were measured in ImageJ and normalized to spheroid area.

### Bulk RNA sequencing and transcriptome data analysis

To characterize the transcriptomic profiles of LNCaP cells under different culture conditions, cells were seeded as 2D monolayers (5,000 cells cm^−2^) or 3D spheroids (3,000 cells per well). and cultured at 21% O_2_ for 3 days. Subsequently, cultures were maintained under normoxic (21% O_2_) or hyperoxic (40% O_2_) conditions for an additional 5 days. Total RNA was extracted and purified by Direct-zol RNA Miniprep Plus Kits (Zymo Research, Cat# R2072) according to the manufacturer’s instructions. The extracted RNA samples were sent to Novogene Corporation Inc. (Sacramento, CA) for quality control, library construction, and sequencing. Briefly, RNA degradation and contamination were monitored on 1% agarose gels. RNA purity was checked using the NanoPhotometer® spectrophotometer (IMPLEN, CA, USA). RNA integrity and concentration were quantified using the RNA Nano 6000 Assay Kit of the Bioanalyzer 2100 system (Agilent Technologies). High-quality RNA samples that passed QC were used to construct sequencing libraries using the NEBNext® Ultra™ RNA Library Prep Kit for Illumina® (NEB, USA) with poly-T oligo-attached magnetic beads for poly-A enrichment. The libraries were sequenced on an Illumina NovaSeq 6000 platform to generate 150-bp paired-end reads at a depth of 20 million reads per sample.

For data analysis, raw reads were processed using FastQC and Trim Galore to remove adapters, low-quality bases (*Q <* 25), and the first 10 bp of the 5’ end to eliminate potential bias. The cleaned reads were aligned to the human reference genome, GRCh38, using STAR, and gene counts were normalized to counts per million (CPM) and analyzed for differential expression using the limma R package. Gene Set Enrichment Analysis (GSEA) was performed using the WebGestalt (WEB-based GEne SeT AnaLysis Toolkit)^48^ web server. Genes were ranked by their t-statistics derived from the limma analysis, and enrichment was assessed against the Reactome pathway database. Pathways with a False Discovery Rate (FDR) *<* 0.05 were considered significantly enriched.

To investigate the transcriptional changes associated with BBR-Enza synergy, compact 22RV1 spheroids (48 h post-seeding) were then treated with vehicle, 20 *µ*M Enza, 5 *µ*M BBR, or the combination under 40% O_2_ for 3 days. Total RNA was extracted and purified by Direct-zol RNA Miniprep Plus Kits according to the manufacturer’s instructions. The extracted RNA samples were submitted to the UCLA Technology Center for Genomics & Bioinformatics (TCGB). Prior to library preparation, RNA integrity was assessed using an Agilent TapeStation system, with all samples exhibiting an RNA Integrity Number *>* 7. Sequencing libraries were generated using the ABclonal FAST mRNA Library Prep Kit with poly-A selection. The libraries were sequenced on an Illumina NovaSeq X Plus platform to generate 50-bp paired-end reads at a depth of 35 million reads per sample. For data analysis, sequencing data quality was confirmed with *>* 90% of bases achieving a Q30 score. High-quality reads were aligned to GRCh38 using STAR. Gene counts were normalized to CPM and analyzed for differential expression using limma R package. GSEA was performed using the GSEA software (Broad Institute). Genes were ranked using the log2 ratio of classes metric based on normalized counts (CPM), and enrichment was assessed against the Hallmark gene set collection (MSigDB) using 2,000 gene set permutations. Pathways with a nominal *p*-value *<* 0.05 and an FDR *<* 0.25 were considered significantly enriched.

### Combo signature gene set identification and function enrichment analysis

To identify a transcriptomic signature specific to the synergistic interaction between BBR and Enza, we employed a subtractive filtering strategy adapted from Fuertes et al.^49^. This refined approach was designed to rigorously remove monotherapy-driven gene expression changes and retain only those genes uniquely associated with the combination response. For all comparisons, statistical significance was defined as an adjusted *P*-value *<* 0.05, and genes that met this threshold were identified as differentially expressed genes (DEGs). To exclude the independent effects of BBR (List 1), we first selected DEGs from the “Combination vs. Enza” comparison (5,662 genes). From this list, we excluded genes that were significantly changed in the same direction (based on the sign of log2 fold change) in the “BBR vs. Control” comparison. Genes that were either non-significant or significantly regulated in the opposite direction in the BBR monotherapy were retained to capture newly induced effects or expression reversal, resulting in 5,258 genes. Similarly, to exclude the independent effects of Enza (List 2), we selected DEGs from the “Combination vs. BBR” comparison (1,667 genes) and excluded genes that were significantly changed in the same direction in the “Enza vs. Control” comparison, retaining those that were non-significant or significantly regulated in the opposite direction under Enza monotherapy (673 genes). Finally, the intersection of List 1 and List 2 yielded 525 genes, comprising 354 up-regulated and 171 down-regulated genes, which were designated as the “Combo Signature.”

To verify the synergistic expression patterns of the identified “Combo Signature”, the expression fold changes of the Combo Signature genes were compared against the monotherapies and their expected additive effect, which was defined using the formula: Log2FC_Additive_ = Log2FC_Enza_ + Log2FC_BBR_. Statistical differences across these four conditions were assessed using a Friedman test, followed by post-hoc pairwise comparisons using paired Wilcoxon signed-rank tests with Holm correction. Subsequently, to elucidate the biological functions of the signature, the 354 up-regulated and 171 down-regulated genes were analyzed separately using g:Profiler^50^. Enrichment was assessed against the Gene Ontology Biological Process (GO:BP) database using the g:SCS significance threshold of *<* 0.05, and representative enriched pathways are presented in Fig. 6h.

### 13C-glucose isotope tracing and metabolomic analysis

Formed LNCaP spheroids (48 h post-seeding) were cultured under 21% or 40% O_2_ for 72 h. To trace glucose metabolism, the complete culture medium was replaced with glucose-free RPMI medium (Gibco, Cat# 11879020) containing 10% dialyzed FBS (Gibco, Cat# A3382001) and 2 g L^−1^ uniformly labeled D-[U-^13^C_6_] glucose (Cambridge Isotope Laboratories, Cat# CLM-1396-1) for the final 24 h of the incubation period. For each condition, 64 spheroids were collected and washed with ice-cold 150 mM ammonium acetate buffer. Intracellular metabolites were extracted with ice-cold 80% methanol containing 0.1 mM norvaline (Sigma-Aldrich, Cat# N7502-25G) as an internal standard. Spheroids were homogenized by brief vortexing, followed by incubation at −80^◦^C for 30 min to facilitate extraction. Lysates were centrifuged at 16,000 × g for 5 min at 4^◦^C, and supernatants were collected and dried under nitrogen using an EZ-2 Elite centrifugal evaporator (Genevac). Cell pellets were collected, and genomic DNA was purified using the GeneJET Genomic DNA Purification Kit (Thermo Scientific, Cat# K0721). Subsequently, DNA concentration was quantified using a NanoDrop 2000C spectrophotometer (Thermo Scientific). Total DNA content was calculated and used to normalize metabolite abundances for downstream comparative analyses.

Dried metabolite extracts were sent to the UCLA Metabolomics Center for LC–MS analysis. Briefly, dried metabolites were reconstituted in 100 *µ*L of 50% acetonitrile (ACN) in water, and 5 *µ*L was loaded onto a Luna NH_2_ column (3 *µ*m, 100 Å, 150 × 2.0 mm; Phenomenex). Chromatographic separation was performed on a Vanquish Flex UHPLC system (Thermo Scientific) using mobile phase A (5 mM ammonium acetate, pH 9.9) and mobile phase B (ACN) at a flow rate of 200 *µ*L min^−1^. A linear gradient from 15% A to 95% A over 18 min was followed by 7 min isocratic flow at 95% A and re-equilibration to 15% A. Metabolites were detected on a Q Exactive Orbitrap mass spectrometer (Thermo Scientific) operated in polarity-switching, full-scan mode over an m/z range of 70–975 with a resolution of 70,000. MAVEN (v8.1.27.11) was used to quantify targeted metabolites by peak area (AreaTop), based on accurate mass measurements (*<* 5 ppm) and previously verified retention times. ^13^C natural abundance correction was performed using AccuCor. Relative amounts of metabolites were calculated by summing up the intensities of all detected isotopologues of a given metabolite. Downstream data processing and statistical analyses were performed using in-house R scripts.

### PSA measurement

Following 72 h incubation at their respective oxygen levels, conditioned media were collected from LNCaP spheroids. PSA concentrations were quantified using the Human Kallikrein 3/PSA Quantikine ELISA Kit (R&D Systems, Cat# DKK300) according to the manufacturer’s instructions. DNA from spheroids was extracted with the GeneJET Genomic DNA Purification Kit and quantified on a NanoDrop 2000C spectrophotometer. PSA values were then normalized to total DNA content per replicate.

### Intracellular reactive oxidative species (ROS) measurement

Following 72 h incubation at their respective oxygen levels, LNCaP spheroids were washed with PBS and incubated with 10 *µ*M H_2_DCFDA (Invitrogen, Cat# D399), a fluorescent probe for intracellular ROS, in PBS at 37^◦^C for 1 h under their respective oxygen conditions. Following incubation, spheroids were washed twice with serum-free medium, and fluorescence intensity (Ex/Em: 495/528 nm) was quantified using a Synergy H1 microplate reader (BioTek Instruments). ROS levels were normalized to the total DNA content of each replicate.

### Dose-response analysis, small-molecule screening, and cell-viability assays

For Enza dose–response assays, PCa spheroids (48 h post-seeding) were treated with Enza at concentrations ranging from 512 *µ*M to 2 *µ*M (2-fold serial dilutions). Vehicle controls were treated with matched concentrations of DMSO. The spheroids were incubated under 21% or 40% O_2_ for 72 h. Absolute cell viability was quantified by live/dead staining and expressed as the ratio of live cells to total cells (live ÷ [live + dead]). Half-maximal inhibitory concentration (IC_50_) values were determined by fitting dose-response data to a four-parameter logistic non-linear regression model using GraphPad Prism software.

Small-molecule screening and validation were performed in a tiered approach. (1) Primary screen: 22RV1 spheroids were treated with 95 MAESTRO-prioritized PKIs (Molecular Screening Shared Resource, UCLA) or 150 NPs (TargetMol) at fixed concentrations (1 or 10 *µ*M), either as single agents or in combination with 20 *µ*M Enza. (2) Multi-dose response validation: The top 14 NP hits were re-evaluated in 22RV1 spheroids at 1–20 *µ*M in the presence of 20 *µ*M Enza. (3) Cross-model validation: Four lead candidates (berberine hydrogen sulphate, nimbolide, deguelin, and resibufogenin) were tested in MDVR, LNCaP, and VCaP spheroids (1–20 *µ*M NPs) combined with model-specific Enza concentrations (20 *µ*M for MDVR, 10 *µ*M for LNCaP, and 20 nM for VCaP). (4) Synergy matrix: A full checkerboard validation was performed for berberine hydrogen sulphate in 22RV1 spheroids, assessing combinations of berberine hydrogen sulphate (0.5–32 *µ*M) and Enza (6.67–180 *µ*M). Across all stages, spheroids (48 h post-seeding) were incubated with drugs under 40% O_2_ for 72 h. Cell viability was quantified using the CellTiter-Glo 3D assay (Promega, Cat# G9683) and normalized to untreated controls.

### Synergy score identification

Inhibition ratio (E, %) was calculated as: *E* = 1 −cell viability (%). Bliss synergy scores were then computed using: *E_ab_* −(*E_a_* + *E_b_* − *E_a_* × *E_b_*), where *E_a_* and *E_b_* are the inhibition ratios for the monotherapy, and *E_ab_*is the inhibition ratio for the combination therapy^51^. Compounds with *E_ab_* greater than *E*_Enza_ and a Bliss score *>* 0.1 were designated as hits and prioritized for subsequent multi-dose validation. The Bliss scores for the full dose–response matrix of the BBR–Enza combination was computed in SynergyFinder^52^ and visualized as a 3D surface plot.

### Statistical analysis

Unless otherwise stated, quantitative data are presented as mean ± standard deviation (SD), with *n* denoting biologically independent replicates as indicated in the figure legends. Statistical analyses were performed using GraphPad Prism (v9.0; GraphPad Software) or R (v4.3.0). For two-group comparisons, statistical significance was determined using unpaired two-tailed Student’s *t*-tests. For comparisons involving more than two groups, one-way Analysis of Variance (ANOVA) was performed, followed by Dunnett’s post-hoc tests for multiple comparisons. To verify the synergistic expression patterns of the identified “Combo Signature”, the Friedman test followed by paired Wilcoxon signed-rank tests with Holm correction was utilized. For RNA-seq differential expression, statistical testing and multiple-testing correction were performed as described in the corresponding section. Unless otherwise stated, a *P*-value *<* 0.05 was considered statistically significant.

### Data availability

The raw and processed RNA sequencing data generated in this study have been deposited in the Gene Expression Omnibus (GEO) database under accession code [GSEXXXXX]. The metabolomics data for 3D prostate cancer spheroids have been deposited in the [MetaboLights/Metabolomics Workbench] database under accession code [IDXXXXX]. The drug screening datasets generated and analyzed during the current study—including the MAESTRO-predicted rankings, aleatoric uncertainty metrics, and experimental Bliss synergy scores for the 1,250 protein kinase inhibitors and 3,952 natural products are provided in the Supplementary Data files. Publicly available datasets used for benchmarking and knowledge-augmentation were accessed from the DrugComb database (v1.4) at https://drugcomb.fimm.fi and PubTator3 at https://www.ncbi.nlm.nih.gov/research/pubtator3/. Any other data supporting the findings of this study are available from the corresponding authors upon reasonable request.

### Code availability

The custom code for the MAESTRO framework has been archived on Zenodo under the DOI [Zenodo DOI 10.5281/zenodo.18361932]. This repository includes the computational scripts for the RAG workflow used for drug-MoA labeling and disease-specific context generation. It also contains the ensemble-consensus ranking algorithms and the GBT surrogate model used for SHAP interpretability analysis.

## Results

### Ensemble Consensus and Mechanistic Reasoning Enable High-Confidence Synergy Prediction

The MAESTRO framework (Fig. 1a) establishes a mechanistically grounded discovery pipeline that anchors LLM reasoning in curated biological priors. Central to this approach is a Retrieval-Augmented Generation (RAG) architecture that systematically synthesizes two molecularly-annotated data streams into context-augmented prompts: (i) compound-specific profiles detailing primary and secondary mechanisms of action (MoA), and (ii) a high-density map of the disease-specific landscape. This context is structured to capture essential biological dimensions, including AR signaling crosstalk, acquired resistance mechanisms, and the upstream regulatory networks that drive adaptive survival. By integrating these disparate mechanistic features, MAESTRO provides the LLM with the domain-specific grounding necessary to prioritize therapeutic candidates based on their predicted capacity for orthogonal inhibition of bypass pathways. This architecture effectively transforms the discovery process from a correlational search into a mechanistically driven rank-ordering of potential synergies.

To translate these mechanistically grounded insights into reproducible quantitative metrics, we implemented an ensemble-ranking strategy that evaluates candidate synergies through batch-wise competitive analysis. Under this framework, the ranking LLM evaluated randomized cohorts of 50 compounds to determine their relative synergy with enzalutamide. For each candidate, the model provided a specific mechanistic justification to support the assigned rank (Fig. S1a). To address the stochastic nature of LLM reasoning and minimize batch-specific bias, we employed an iterative sampling strategy across diverse compound cohorts. This approach stabilized the ranking process, allowing mean ranks to achieve convergence with less than 1% variation after 100 independent runs (Fig. S1b). The resulting architecture yields a dual-metric output consisting of a consensus mean rank, which represents predicted synergistic potential, and a rank standard deviation that serves as a quantitative measure of aleatoric uncertainty. Benchmarking against 150 AR inhibitor-anchored compound pairs from the DrugComb database^53^ confirmed the predictive power of this ensemble-ranking framework. This analysis confirmed the predictive power of the ensemble-ranking framework and revealed a distinct "inverted U-shape" relationship between mean rank and rank standard deviation (SD) (Fig. 1b). We observed that the LLM achieves maximum internal consensus at the ranking extremes of high synergy and antagonism. Motivated by this observation, we defined the lowest-variance quartile as the "High-Confidence Zone" (HCZ), where the model’s prediction is most consistent. Within this HCZ, the Spearman correlation between predicted and experimental rankings reached *ρ* ∼ 0.87 (Fig. 1c–d), whereas expanding the inclusion threshold to high-variance candidates caused the correlation to degrade toward 0.1. This predictive accuracy trend and the underlying mean-variance relationship were consistently reproduced across independent abiraterone-anchored datasets (Fig. S1c–d) and alternative architectures, including Google Gemini (Fig. S1e–f). Collectively, these results demonstrate that ensemble consensus serves as a robust proxy for prediction accuracy, allowing MAESTRO to autonomously identify its most reliable therapeutic leads.

Beyond benchmarking accuracy, we interrogated the architectural dependencies and translational utility of MAESTRO to identify the essential drivers of the discovery process. Evaluating the specific contribution of the RAG module revealed that mechanistic knowledge is the primary driver of predictive accuracy. Removing this augmentation layer caused Spearman correlations within the HCZ to fall from 0.90 to 0.43 (Fig. S1d). Furthermore, the framework exhibited architectural invariance, maintaining high cross-platform consistency between ChatGPT and Gemini backends (*ρ* ∼0.97) specifically within the HCZ (Fig. S1g–h). Finally, to operationalize these findings for high-throughput discovery, we quantified the impact of stringent ranking and variance thresholds on predictive precision. Restricting selection to the top 33% of candidates within the lowest-variance window yielded a precision of approximately 70%, representing a two-fold enrichment over the random baseline (Fig. 1e). These optimization data suggest that the MAESTRO architecture can effectively concentrate experimental resources on the highest-probability therapeutic leads, thereby minimizing the screening burden associated with false-positive results.

### MAESTRO Identifies Novel Synergistic Combinations through Mechanistically Grounded Inference

We next tested the hypothesis that MAESTRO could resolve previously unrecognized synergies by synthesizing latent molecular relationships within its curated knowledge base. To evaluate this discovery potential, we applied the framework to a library of 1,250 protein kinase inhibitors (PKIs) to prioritize compounds for combination with enzalutamide. This library provided a pharmacologically well-defined substrate to assess whether the model’s integration of signaling crosstalk could extend beyond the recapitulation of known drug pairs. Virtual screening successfully reproduced the characteristic "inverted U-shaped" uncertainty distribution (Fig. 2a), allowing us to prioritize 95 lead candidates within the HCZ.

**Figure 2.**
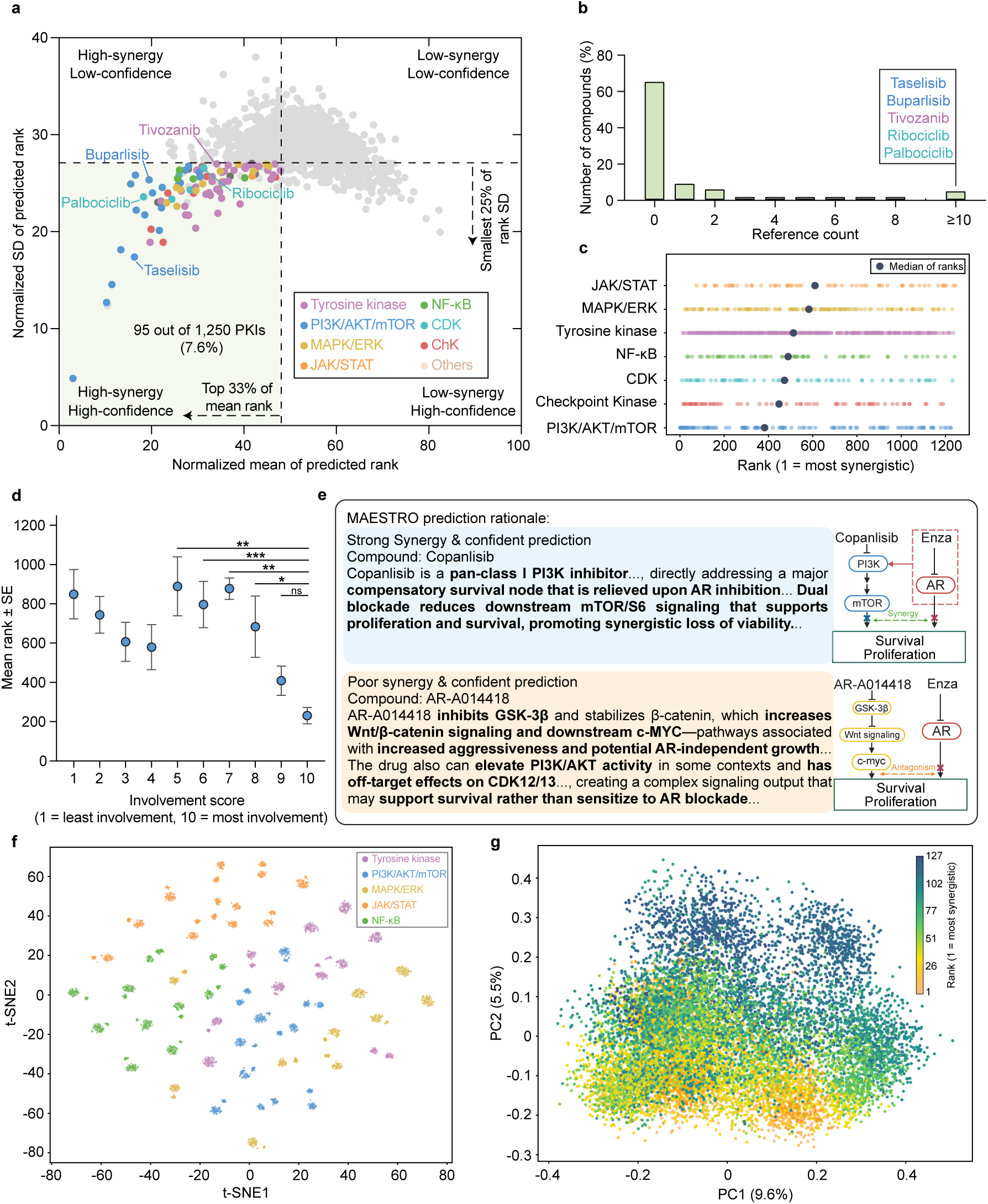
MAESTRO demonstrates interpretable mechanistic reasoning through kinase inhibitor prioritization. **(a)** Scatter plot of MAESTRO-predicted normalized standard deviation of predicted rank versus normalized mean of predicted rank for 1,250 PKIs (400 runs per compound). Selection of 95 prioritized candidates was defined by a top 33% cutoff for mean rank and a smallest 25% cutoff for rank SD. **(b)** Frequency of publications reporting synergy between enzalutamide and each of the 95 lead compounds. Data were obtained via manual curation of Google Scholar. **(c)** Distribution of predicted synergy ranks across seven mechanism-based categories, with median ranks indicated by a black dot for each category. Low rank value corresponds to high synergy prediction. **(d)** Mean predicted synergy rank plotted against involvement score for 127 inhibitors within the PI3K/AKT/mTOR pathway. This LLM-derived score (1–10) quantifies the directness of targeting key pathway nodes, where higher values signify core pathway inhibition. Data are presented as mean ± SE. **(e)** Representative MAESTRO-generated prediction rationales accompanied by schematic illustrations comparing high-(blue) and low-synergy (yellow) candidates within the PI3K/AKT/mTOR pathway category. The red box in the inset highlights AR inhibition, which triggers a compensatory signaling pathway**. (f)** t-SNE visualization of semantic embeddings derived from generated rationales across five categories (10 compounds per category; 100 rationales per compound). **(g)** PCA plot of rationale embeddings for the PI3K/AKT/mTOR-associated category (127 compounds; 100 rationales per compound). Data points are colored by predicted synergy rank. Statistical significance was determined by one-way ANOVA followed by Dunnett’s multiple comparisons test; Statistical significance is denoted by asterisks, where * indicates p < 0.05, ** indicates p < 0.01, *** indicates p < 0.001, and ns indicates no statistically significant difference.

While tyrosine kinase inhibitors were numerically dominant in the library, inhibitors targeting the PI3K/AKT/mTOR axis exhibited the highest enrichment rate (15.7%; Fig. S2a). Crucially, a systematic literature cross-validation of these 95 prioritized leads revealed that over 65% lacked any prior reports of synergy with enzalutamide (Fig. 2b). These findings suggest that by mapping compound-specific mechanisms onto the broader landscape of resistance signaling, MAESTRO can autonomously predict novel synergistic interactions that remain latent in the existing biomedical corpus.

To understand the biological logic underpinning these predictions, we first characterized the macro-level features driving MAESTRO’s decision-making. Inter-pathway analysis of the 1,250 PKIs revealed noticeable shifts in median rank distributions across categories, confirming that the model incorporates pathway-level selectivity into its predictive logic (Fig. 2c). Moreover, the presence of significant intra-pathway variance suggests that MAESTRO distinguishes between individual molecular profiles within the same functional class. To dissect the sources of this intra-group variation, we analyzed target involvement within the top-ranked PI3K/AKT/mTOR pathway to characterize the logic driving its specific rankings. Specifically, an LLM-derived involvement score (1–10) was applied to each of the 127 inhibitors to quantify their directness in targeting key pathway nodes. We found that these involvement scores correlated with MAESTRO rankings, where higher scores, indicating proximity to the pathway core, mirrored more favorable predictive priorities (Fig. 2d).

This logic is exemplified by the contrasting prioritizations of Copanlisib and AR-A014418 (Fig. 2e). Copanlisib, identified as a core PI3K/AKT/mTOR axis inhibitor with a high involvement score, was highly ranked for its capacity to block compensatory survival signaling. Conversely, AR-A014418 received a low score and low prioritization due to its targeting of peripheral nodes and potential for antagonism. MAESTRO’s mechanistic justifications distinguished these cases, prioritizing core pathway disruption over the less relevant targeting of distal nodes.

To validate the alignment between numerical rankings and model justifications, we performed embedding-based semantic analysis of the LLM-generated rationales. t-SNE visualization revealed that rationales for distinct compound classes formed well-defined semantic clusters, indicating that the model’s reasoning is mechanism-conditioned (Fig. 2f). Moving from pathway-level to intra-pathway analysis, Principal Component Analysis (PCA) of PI3K/AKT/mTOR rationales revealed a semantic gradient along the PC2 axis that aligned with numerical rankings (Fig. 2g). This relationship was substantiated by word-cloud analysis (Fig. S2b), which showed that high-priority rationales were enriched with terms describing core pathway modulation (e.g., mTOR,” dual blockade,” “compensatory”), whereas low-ranked compounds were associated with peripheral targets. Collectively, these findings confirm that MAESTRO leverages a context-aware mechanistic framework to identify high-confidence candidates and provide reasoning.

### Oxygen-Supplemented 3D Prostate Cancer Spheroids Provide a Physiologically Relevant Validation Platform

MAESTRO substantially narrowed the candidate drug space. This efficiency allowed us to utilize 3D prostate cancer spheroids to validate the nominated drug pairs (Fig. 3a). While 3D models offer superior physiological relevance compared to 2D cultures, they are historically difficult to scale and prone to limited oxygen diffusion. To address these hurdles, we developed an oxygen-supplemented platform that is compatible with standard 96/384-well formats, providing a scalable, high-throughput system that eliminates the necrotic cores typically found in conventional spheroids. Through systematic optimization, we identified 40% O_2_ as the ideal concentration to enhance core penetration for 500–600 *µ*m spheroids (Fig. S3a–c). Staining for Live/Dead, Ki67/CC3, and E-cadherin confirmed that this oxygen supplementation eliminated necrotic artifacts while preserving spheroid viability, proliferation, and structural integrity (Fig. 3b–d, S3d–e, S4a–c). Furthermore, *γ*-H2AX staining, along with ROS and PSA levels, remained comparable to controls at 40% O_2_, confirming the absence of significant oxidative stress or functional impairment (Figs. 3e–f, S4d–f).

**Figure 3.**
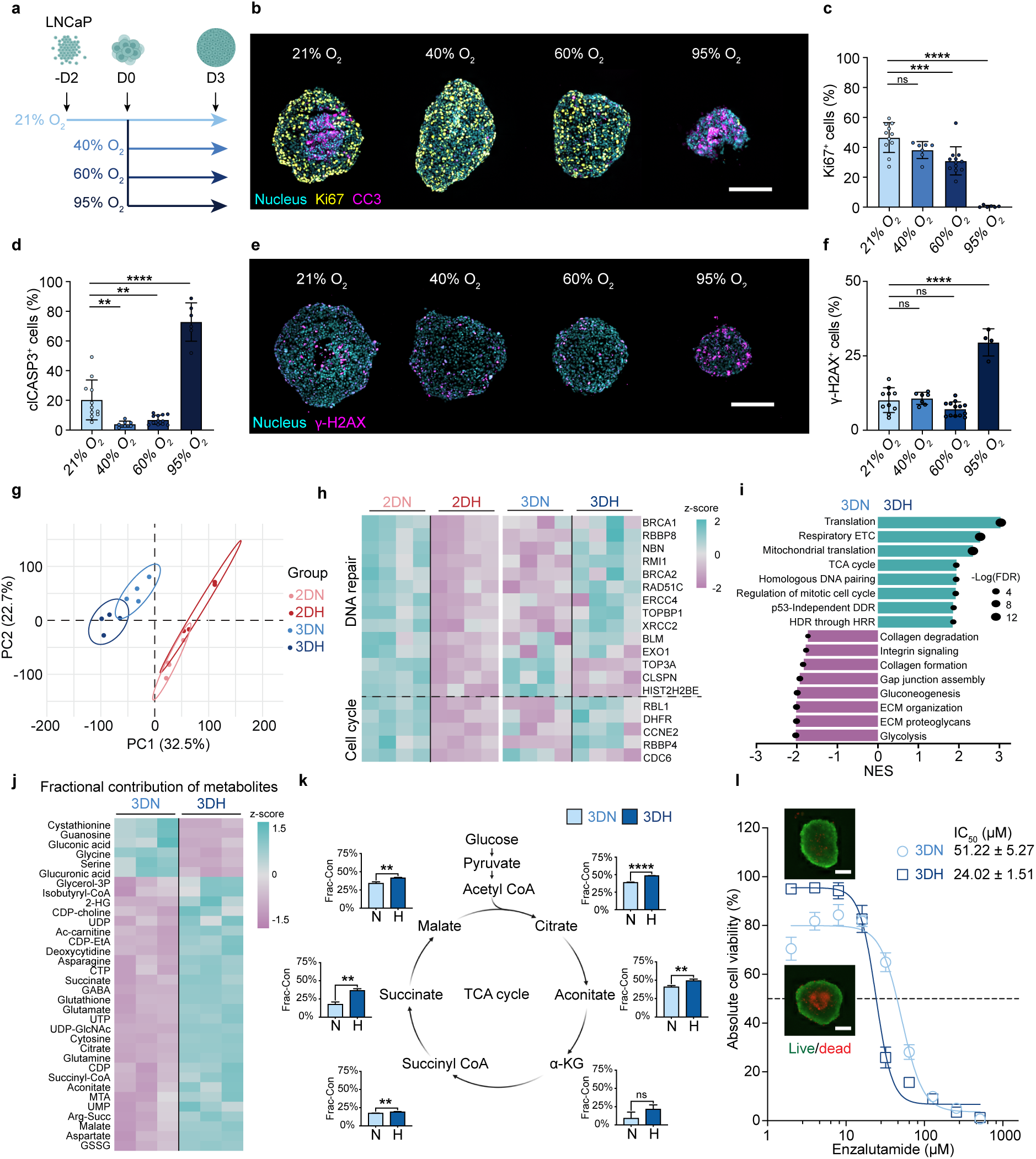
Oxygen-supplemented 3D prostate cancer spheroids provide a high-fidelity validation platform. **(a)** Schematic illustration of the oxygen supplementation experiment. LNCaP spheroids were cultured under varying oxygen levels (21–95% O_2_) to identify the optimal condition for minimal necrotic core formation. **(b)** Representative immunofluorescence images of cells stained for the proliferation marker Ki67 and the apoptosis marker cleaved caspase-3 (CC3) following 3 days of oxygen-supplemented culture. Scale bar, 200 *µ*m. **(c-d)** Quantification of percentage of (**c**) Ki67^+^ cells and (**d**) CC3^+^ cells under the indicated oxygen levels (*n* = 6–12 per group). **(e)** Representative immunofluorescence images of *γ*-H2AX (DNA damage marker) following 3 days of oxygen-supplemented culture. Scale bar, 200 *µ*m. **(f)** Quantification of percentage of *γ*-H2AX^+^ cells under indicated oxygen levels (*n* = 4–13 per group). **(g)** PCA plot of RNA-seq profiles across four culture conditions: 2D normoxia (2DN; 21% O_2_), 2D hyperoxia (2DH; 40% O_2_), 3D normoxia (3DN), and 3D hyperoxia (3DH). Ellipses represent 95% confidence ellipses. **(h)** Heatmap visualizing the z-scored expression patterns of DNA repair and cell cycle-related genes across the four groups. **(i)** Gene set enrichment analysis comparing 3DH versus 3DN. The bar chart displays representative upregulated and downregulated pathways, determined by the normalized enrichment scores (NES). The size of the black dot represents the −log(FDR), where FDR denotes the false discovery rate. Data in **(g–i)** are derived from *n* = 4 biological replicates. **(j)** Heatmap of z-scored fractional contributions of ^13^C-glucose labeled metabolites in 3DH and 3DN conditions. **(k)** Bar charts depicting fractional contributions of TCA cycle intermediates comparing 3DH and 3DN. The middle schematic illustrates the TCA cycle. Data in **(j–k)** are derived from *n* = 3 biological replicates. **(l)** Enzalutamide dose-response curves plotting absolute viability derived from Live/Dead staining for LNCaP spheroids in 3DN and 3DH groups (*n* = 4). Representative images of untreated controls at Day 3 are shown as insets. Scale bar, 200 *µ*m. Data are presented as mean ± SD. Statistical significance was determined by one-way ANOVA followed by Dunnett’s multiple comparisons test (**c, d, f**) or two-tailed unpaired Student’s t-test (**k**); * *P <* 0.05, ** *P <* 0.01, *** *P <* 0.001, **** *P <* 0.0001, ns: not significant.

Omics analyses confirmed that the 3D environment protects against hyperoxic stress; transcriptomic profiles showed that the DNA repair and cell cycle suppression seen in 2D hyperoxia were reversed in our spheroids (Fig. 3g–i, Fig. S5a–c). This lack of genomic stress was paired with a highly productive metabolism. Specifically, gene set enrichment analysis (GSEA) and metabolomic data revealed that 40% O_2_ enhanced glucose incorporation into the TCA cycle, supporting a metabolically active and proliferative phenotype (Fig. 3i–k, Fig. S6a–b). We also evaluated whether oxygenated spheroids can resolve necrosis-driven screening limitations by performing enzalutamide dose-response assays. At 40% O_2_, spheroids maintained near 100% baseline viability and showed heightened sensitivity to low-dose enzalutamide. In contrast, 21% O_2_ cultures were compromised by necrosis-associated masking, yielding only ∼75% viability in untreated and low-dose groups (Fig. 3l, Fig. S5d). By eliminating necrotic cores while preserving high viability and physiological relevance, it provides a robust system for validating MAESTRO-predicted candidates.

### *In vitro* Validation Confirms MAESTRO’s High Hit Rates and Pathway-level Accuracy

Using oxygenated 3D spheroids of enzalutamide-resistant 22RV1 and C4-2B MDVR (MDVR)^46^ cell lines, we validated the synergy of 95 MAESTRO-prioritized PKIs. After 72 hours of combination treatment (1 *µ*M PKI and 20 *µ*M enzalutamide), we assessed cell viability (Fig. 4a) and calculated synergy using Bliss scores^51^. Following standard convention, we defined “hits” as compounds exhibiting a Bliss score > 0.1 and a combinatorial efficacy (E_ab_) exceeding enzalutamide monotherapy. In 22RV1 spheroids, we identified 17 hits, a 17.9% hit rate that is ∼100-fold higher than the typical yield of conventional exhaustive screenings (∼0.05%)^54,55^. Among the screened compounds, PI3K/AKT/mTOR inhibitors emerged as a consistently synergistic class with enzalutamide in both 22RV1 and MDVR spheroids (Fig. 4b and Fig. S7a).

**Figure 4.**
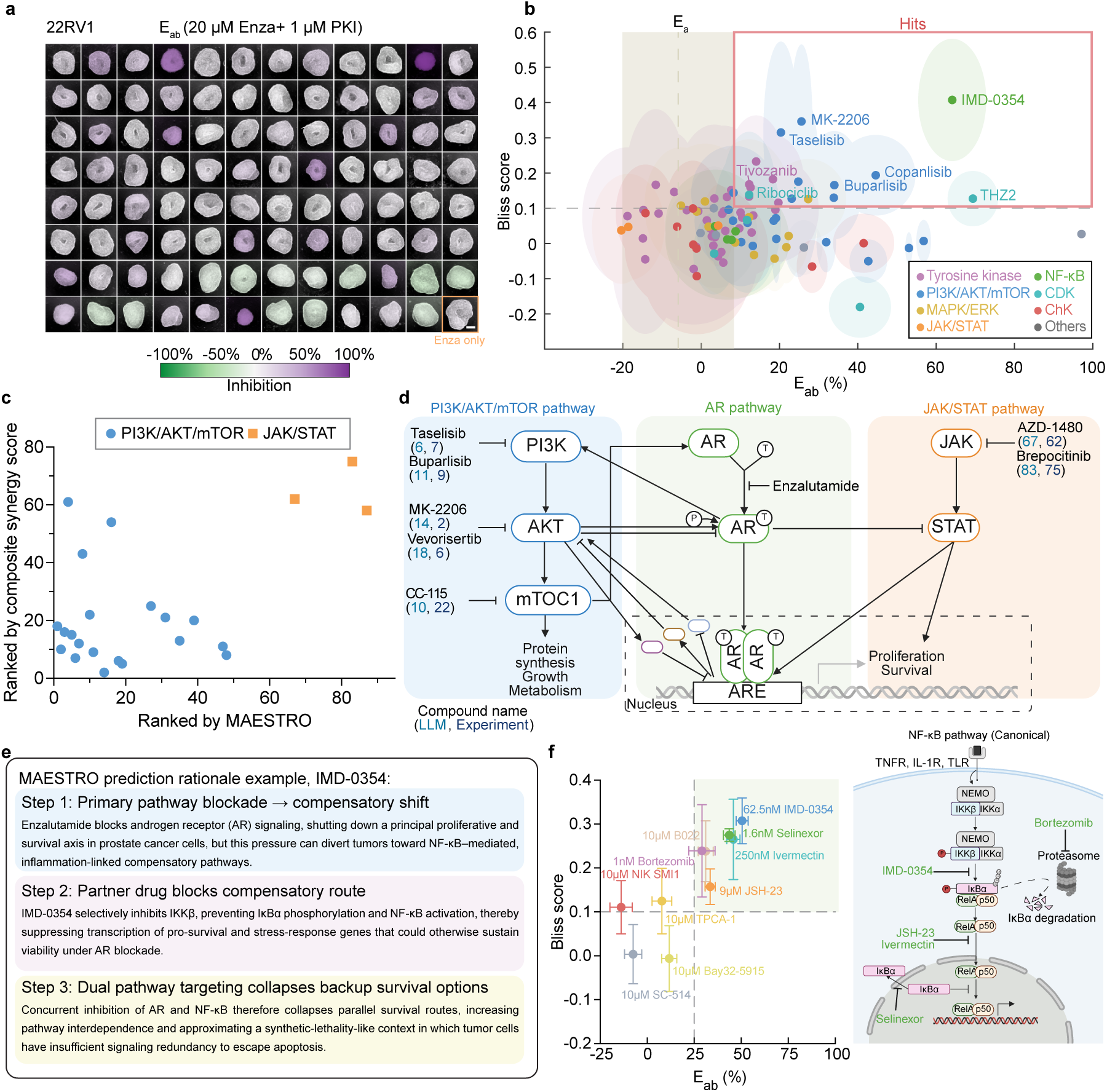
Experimental validation confirms MAESTRO’s predictive accuracy and pathway-level reasoning in enzalutamide-resistant models. **(a)** Representative pseudocolored bright-field images of oxygen-supplemented 3D 22RV1 spheroids treated with 95 MAESTRO-prioritized PKIs in combination with enzalutamide for 3 days under 40% O_2_. Images are pseudocolored according to the combinatorial inhibition ratio (E_ab_) relative to untreated controls. The enzalutamide monotherapy group (bottom left, orange box) serves as the therapeutic benchmark for synergy evaluation. Scale bar, 200 *µ*m. **(b)** Scatter plot of E_ab_ versus Bliss scores for the 95 PKIs tested in 22RV1 spheroids. E_ab_ was calculated as the inhibition rate (%) relative to the untreated group. Hits (red boxed region) are defined as compounds exhibiting E_ab_ greater than enzalutamide monotherapy (E_a_) and a Bliss score *>* 0.1. Error ellipses denote the SD (*n* = 4) for E_ab_ (x-axis) and Bliss scores (y-axis). The vertical dashed line and shaded box represent the mean ± SD of the E_a_. **(c)** Scatter plot comparing MAESTRO-predicted rankings with composite synergy score rankings for PKIs targeting the PI3K/AKT/mTOR and JAK/STAT pathways. The composite synergy score was defined as Bliss_scaled_ × *E_ab,_*_scaled_, where mean Bliss and E_ab_ (inhibition, %) were first averaged across 22RV1 and MDVR spheroids for each PKI and then min–max scaled to 0–1 across all 95 screened PKIs. **(d)** Schematic illustration of signaling crosstalk between PI3K/AKT/mTOR and AR signaling compared to the JAK/STAT and AR signaling. Representative compounds are annotated with their MAESTRO predictions and experimental rankings derived from composite synergy scores. **(e)** MAESTRO-generated mechanistic rationale for the hit compound IMD-0354, illustrating the model’s step-by-step reasoning. **(f)** Pathway validation of the NF-*κ*B signaling axis as a therapeutic target. Left: Scatter plot of E_ab_ versus Bliss scores for 10 NF-*κ*B inhibitors. E_ab_ was calculated as the inhibition rate (%) relative to the untreated group (*n* = 4). Right: Schematic of the canonical NF-*κ*B pathway, mapping the molecular targets of the identified hit compounds. Data are presented as mean ± SD.

Aggregated experimental rankings aligned with the priorities established by the MAESTRO framework (Fig. S7b–c). Notably, this validation was restricted to the top 7.6% (95/1,250 compounds) of prioritized candidates. By excluding the low-performing noise characteristic of unselected libraries, this subset presents a truncated distribution that inherently challenges the resolution of standard correlation metrics. Despite this narrow window, the framework demonstrated discriminatory precision. High-potential PI3K/AKT/mTOR inhibitors clustered prominently in the high-synergy quadrant, whereas JAK/STAT inhibitors remained largely non-responsive (Fig. 4c). These results indicate that the framework’s context-aware reasoning captures the underlying signaling topology of prostate cancer. The model correctly prioritized the PI3K/AKT/mTOR axis, which engages in robust reciprocal feedback with AR signaling to function as a primary adaptive bypass mechanism^6,7,12^. Conversely, the model accurately deprioritized the JAK/STAT axis due to its limited functional connectivity within this lineage (Fig. 4d). These findings demonstrate that by mapping latent signaling dependencies, MAESTRO achieves the mechanistic resolution necessary to differentiate synergies from non-responsive candidates within enriched therapeutic landscapes.

Beyond making accurate pathway-level prioritizations, MAESTRO identified the IKK*β* inhibitor IMD-0354 as a novel hit, a finding we experimentally confirmed across both enzalutamide-resistant cell lines (Fig. S7d). MAESTRO provided a mechanistic rationale for this synergy (Fig. 4e), proposing that AR blockade diverts cancer cells toward inflammation-linked survival pathways that can be disrupted by targeting the NF-*κ*B compensatory node. To evaluate whether this signaling vulnerability represents a conserved therapeutic target, we performed a validation screen using ten structurally diverse NF-*κ*B inhibitors within the 22Rv1 spheroid model. This confirmatory screen yielded a 60% hit rate (Fig. 4f, Fig. S7e), with the majority of hits targeting the canonical NF-*κ*B pathway. These results validate MAESTRO’s rationale that targeting the NF-*κ*B axis is a robust strategy for overcoming enzalutamide resistance.

### MAESTRO Deconvolves Multi-Target Drug Effects and Identifies Berberine as a Synergistic Agent

Having demonstrated MAESTRO’s performance on single-target inhibitors, we next sought to evaluate its ability to resolve the complex multi-target interactions inherent to natural products (NPs). While the broad target profiles of NPs traditionally hinder their development as selective monotherapies, this inherent polypharmacology makes them ideal candidates for combination strategies designed to achieve multi-node disruption of resistant signaling networks^56^. We applied the framework to a library of 3,952 NPs to assess its capacity to navigate these complex mechanistic landscapes across varying levels of prior annotation. As expected, the prioritization yielded a characteristic inverted U-shaped uncertainty distribution (Fig. 5a), identifying a subset of high-priority compounds whose multi-target mechanisms align with the requirements for orthogonal inhibition of AR-bypass pathways.

**Figure 5.**
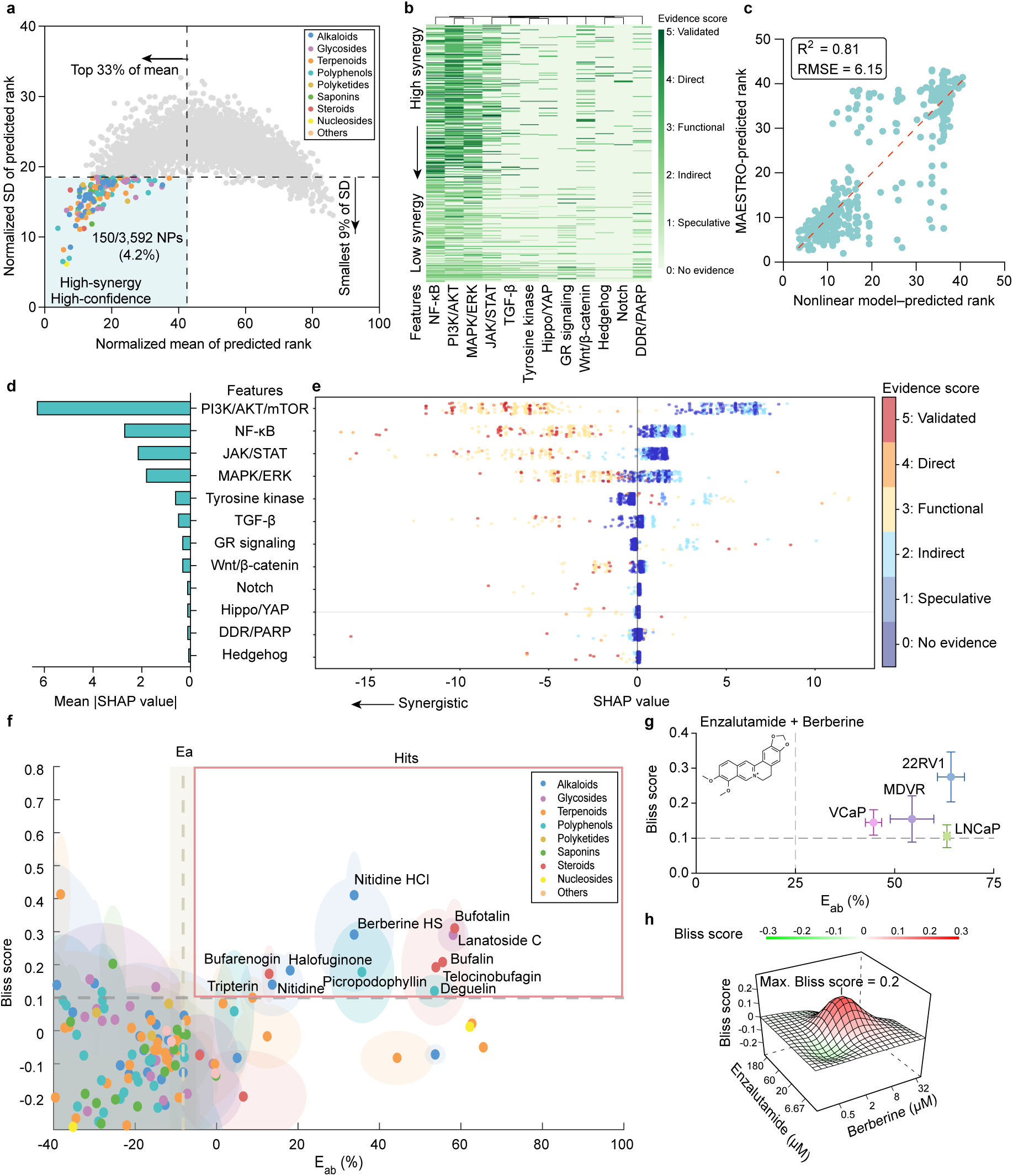
Deciphering MAESTRO’s prioritization logic in polypharmacology and validating synergistic natural product candidates. **(a)** Distribution of 3,592 natural products based on MAESTRO-predicted mean synergy rank and rank SD (200 runs per compound). The highlighted region denotes 150 prioritized candidates for experimental validation, defined by a top 33% cutoff for mean rank and the lowest 9% cutoff for rank SD. **(b)** Heatmap displaying the evidence scores of 12 signaling features against the mean synergy rank for the top 500 compounds with the lowest rank SD. Evidence scores were quantified by the LLM, and the criteria are detailed in the Methods. Columns (features) were arranged by hierarchical clustering using correlation distance and average linkage, while rows (compounds) were ordered by ascending mean rank (top: high synergy; bottom: low synergy). **(c)** Correlation between Gradient Boosting Tree (GBT)-predicted and MAESTRO-predicted ranks for the 500 compounds with the lowest rank SD. GBT predictions were derived from 12-dimensional evidence vectors. The red dashed line indicates the simple linear regression fit. *R*^2^ denotes the coefficient of determination, and RMSE represents the root mean square error between the two ranking models. **(d–e)** Global feature importance **(d)** and feature impact distribution **(e)** derived from SHAP analysis. In **(d)**, features are ranked by mean absolute SHAP value. In **(e)**, points represent the distribution of SHAP values for 500 compounds across each of the 12 features, colored by evidence score, where negative values indicate a drive towards higher predicted synergy. **(f)** Experimental screening of 150 prioritized candidates (10 *µ*M) combined with Enzalutamide (20 *µ*M) in 22RV1 spheroids. Scatter plot shows combination efficacy (E_ab_) versus Bliss scores. E_ab_ was calculated as the inhibition rate (%) relative to the untreated group. Hits (red box): E_ab_ *>* enzalutamide monotherapy (E_a_) and Bliss *>* 0.1. Error ellipses: SD (*n* = 4). Vertical line/shade: mean ± SD of E_a_ (*n* = 20). **(g)** Scatter plot of E_ab_ versus Bliss score Validating the synergy of berberine hydrogen sulphate and enzalutamide across four prostate cancer spheroid models (22RV1, MDVR, VCaP, and LNCaP). Data points represent the most synergistic combination derived from the dose-response matrix for each model. Dashed lines indicate the thresholds of E_ab_ and Bliss scores for identifying hits. Data are presented as mean ± SD (*n* = 8). **(h)** 3D synergy plot of the Berberine–Enzalutamide combination (*n* = 3).

**Figure 6.**
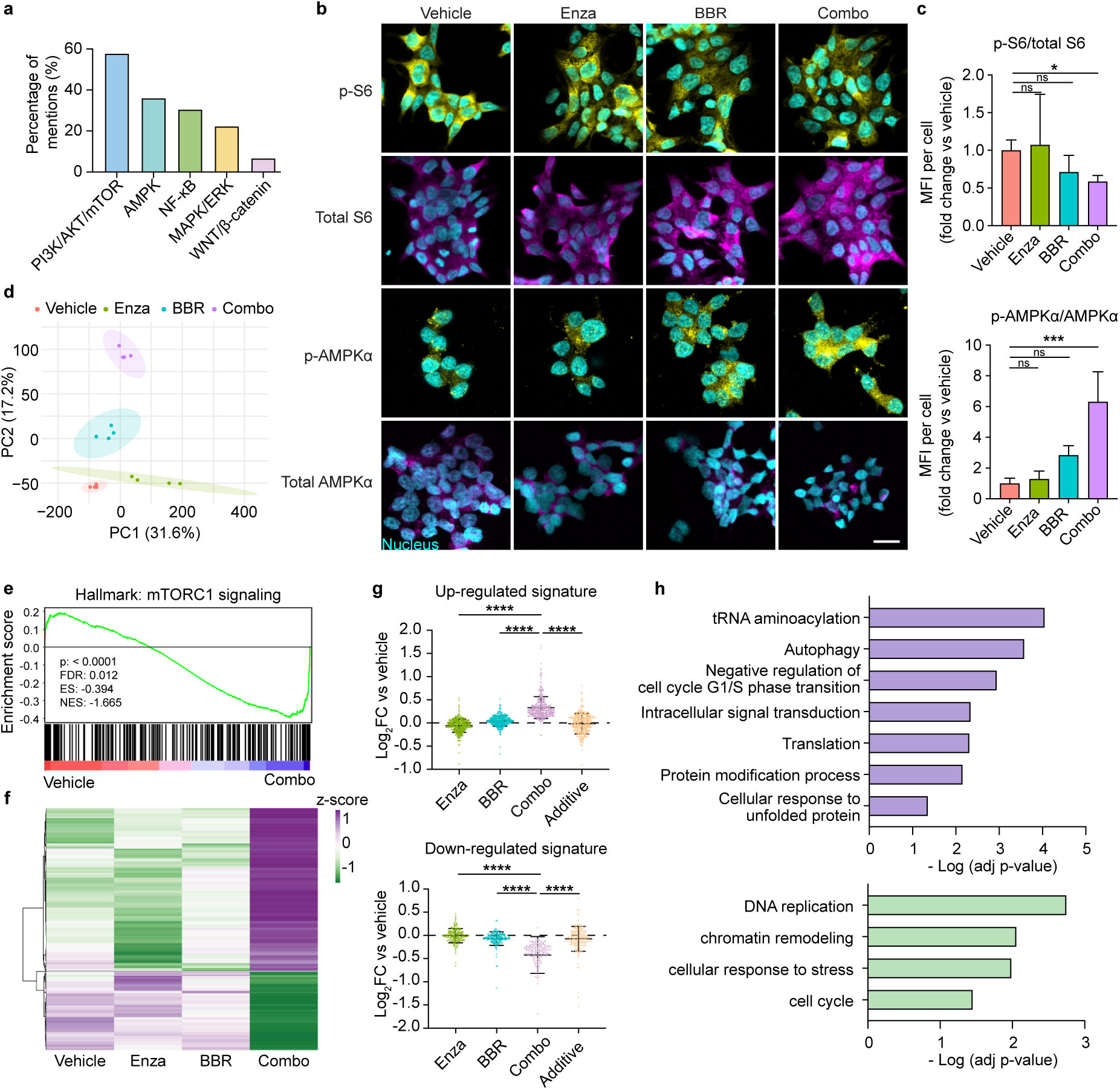
Validation of MAESTRO’s mechanistic justifications for Berberine–Enzalutamide synergy. **(a)** Frequency analysis of MAESTRO-predicted mechanistic rationales contributing to the synergy between berberine and enzalutamide. **(b–c)** Experimental valida-tion of PI3K/AKT/mTOR and AMPK pathway modulation. **(b)** Representative immunofluorescence images and **(c)** corresponding quantification of p-S6/Total S6 (top) and p-AMPK*α*/Total AMPK*α* (bottom) ratios in 22RV1 cells treated with vehicle, 20 *µ*M enzalutamide (Enza), 5 *µ*M berberine (BBR), or the combination (Combo) for 8 h. Scale bar, 20 *µ*m. Data in **(c)** are presented as the fold change of mean fluorescence intensity (MFI) per cell relative to vehicle (*n* = 3, with an average of 613 cells analyzed per replicate). Statistical significance was determined by one-way ANOVA followed by Dunnett’s multiple comparisons test. **(d)** PCA plot visualizing transcriptomic variation (log_2_-CPM) in 3D 22RV1 spheroids across the four treatment groups after 8 h. Ellipses indicate 95% normal probability confidence regions (*n* = 4). **(e)** GSEA plot demonstrating significant downregulation of the Hallmark mTORC1 signaling signature in the Combo group compared to the vehicle group. Normalized enrichment score (NES) and false discovery rate (FDR) are indicated. **(f)** Heatmap displaying the expression profiles of the “Combo Signature,” comprising genes synergistically upregulated (*n* = 354) or downregulated (*n* = 171) upon combination treatment. Rows (genes) were arranged by hierarchical clustering using correlation distance and average linkage. **(g)** Plot depicting expression changes (log_2_FC) of the up-regulated (top) and down-regulated (bottom) signature genes across the treatment conditions relative to the vehicle, compared with the theoretical values for the additive effect of individual therapies. Statistical significance was determined by the Friedman test followed by post hoc pairwise Wilcoxon signed-rank tests with Holm’s correction for multiple comparisons. **(h)** Functional enrichment analysis of the upregulated (top) and downregulated (bottom) Combo Signature genes via g:Profiler. Data are presented as mean ± SD; * *P <* 0.05, ** *P <* 0.01, *** *P <* 0.001, **** *P <* 0.0001, ns: not significant.

While MAESTRO generates granular mechanistic rationales for individual predictions, we sought to characterize the global, systemic logic governing its prioritization of pathway crosstalk at scale. To transform these qualitative justifications into a transparent decision surface, we mapped the inhibition profiles of the natural product library by scoring the likelihood of target engagement across 12 canonical signaling pathways. This high-dimensional analysis enabled us to deconvolve the biological drivers of the discovery process. The resulting evidence scores revealed that MAESTRO preferentially prioritizes distinct inhibitory combinations involving the NF-*κ*B, PI3K/AKT, MAPK/ERK, and JAK/STAT pathways (Fig. 5b). To further dissect how MAESTRO integrates nonlinear signaling crosstalk into its synergy predictions, we utilized a Gradient-Boosted Trees (GBT) surrogate model^44^. This approach allowed us to map compound rankings as a functional output of pathway evidence scores, providing a transparent medium to observe the model’s internal decision logic. The GBT surrogate accurately mimicked MAESTRO’s prediction trend, achieving an *R*^2^=0.81 for the 500 compounds in the low-variance zone (Fig. 5c). The GBT’s prediction accuracy improved further when restricted to the top 100 high-confidence predictions, reaching an *R*^2^=0.93 (Fig. S8a).

To deconstruct the hierarchical signaling logic captured by this GBT surrogate, we utilized SHAP (SHapley Additive exPlanations) analysis^45^ to quantify how specific pathway hubs and their interactions dictate the model’s output. PI3K/AKT/mTOR inhibition emerged as the primary driver of predicted synergy, followed by the NF-*κ*B, JAK/STAT, and MAPK/ERK axes (Fig. 5d). Notably, the SHAP swarm plot (Fig. 5e) reveals noticeable dispersion among PI3K/AKT/mTOR data points with high evidence scores (red and orange dots), with SHAP values ranging from -1 to -12. This horizontal spread for a fixed evidence level indicates that PI3K blockade interacts non-linearly with the broader signaling landscape; its effect is likely potentiated by the concurrent inhibition of secondary pathways, rather than acting in an additive manner. Conversely, importance scores for auxiliary pathways like Notch and Hedgehog remain clustered near zero across all evidence levels, indicating these nodes exert negligible influence on the model’s synergy predictions compared to the core survival hubs.

Lastly, we applied HCZ selection criteria (lowest 9% rank standard deviation; top 33% mean rank) and identified 150 candidates (4.2% of the library) for validation in 3D 22Rv1 spheroids under physiological oxygen supplementation. To account for the typically lower potency of natural products compared to synthetic small molecules, these candidates were screened at 1 and 10 *µ*M in combination with 20 *µ*M enzalutamide. This primary screen yielded 23 unique synergistic hits (17 at 10 *µ*M and 12 at 1 *µ*M, with 6 overlapping), representing a 15.3% hit rate and confirming MAESTRO’s utility in prioritizing bioactive compounds from complex libraries (Fig. 5f and Fig. S8b). Following multi-dose response surface modeling, the leads were further narrowed to four top-tier candidates—berberine, deguelin, nimbolide, and resibufogenin—based on the consistency of their synergistic scores (Fig. S8c). To evaluate generalizability, these candidates were cross-validated across a panel of prostate cancer lines, including enzalutamide-resistant (MDVR) and sensitive (VCaP, LNCaP) models (Fig. S8d). Berberine emerged as the most robust lead, maintaining a stable synergistic profile across all four lines (Fig. 5g). Checkerboard synergy assays (Fig. 5h) further revealed a broad synergistic window at clinically achievable concentrations (enzalutamide: 6–60 *µ*M; berberine: 0.5–8 *µ*M).

### Molecular and transcriptomic profiling validates MAESTRO-predicted mechanisms of pharmacological synergy

While we experimentally confirmed the synergistic potency of the berberine-enzalutamide combination, did the MAESTRO’s underlying justification accurately reflect the true biological dependencies? To address this, we quantified the frequency of pathway targets cited by MAESTRO across 200 independent iterations and identified the dual modulation of PI3K/AKT/mTOR and AMPK signaling as the primary drivers of berberine prioritization (Fig. 6a). We then sought to experimentally validate these computational leads by characterizing the signaling perturbations induced by the combination therapy in 22Rv1 cells via immunofluorescence (Fig. 6b). The phosphorylation of ribosomal protein S6, a canonical downstream effector of mTORC1, was significantly attenuated by the combination therapy, whereas neither monotherapy achieved statistical significance compared to vehicle controls. (Fig. 6c, top). In parallel, the combination treatment markedly upregulated the activation of the energy sensor AMPK relative to vehicle controls, an effect that surpassed both single-agent treatments (Fig. 6c, bottom). Collectively, these findings corroborate the mechanistic accuracy of the predictive justification provided by the MAESTRO framework, confirming that its prioritization is grounded in the targeted disruption of key metabolic and survival nodes.

Next, we performed bulk RNA sequencing on 3D 22RV1 spheroids to obtain a comprehensive view of the downstream cellular responses to the combination treatment. We prioritized this cell line for mechanistic profiling as it served as our primary screening model and exhibited the most robust synergistic phenotype in validation assays. Principal Component Analysis (PCA) illustrated that the combination therapy induced a transcriptional shift along the same trajectory as the monotherapies but with a substantially amplified magnitude (Fig. 6d), suggesting a cooperative enhancement of downstream signaling. Consistent with our immunofluorescence data, Gene Set Enrichment Analysis (GSEA) confirmed the potent downregulation of mTORC1 signaling in the combination group (Fig. 6e).

We then implemented a subtractive transcriptomic filter to isolate the specific mediators of synergy, excluding genes regulated by either monotherapy to enrich for a signature unique to the combination interaction. This approach identified 525 genes (354 up-regulated; 171 down-regulated) whose differential expression significantly exceeded the predicted additive effects of the constituent monotherapies (Fig. 6f–g). Functional enrichment analysis revealed that the up-regulated signature was associated with adaptive stress responses, including autophagy, negative regulation of cell cycle G1/S phase transition, and the unfolded protein response (UPR), which are consistent with an AMPK-activated and mTORC1-inhibited state^57–60^ (Fig. 6h).

Concurrently, the down-regulated signature was significantly enriched in DNA replication and cell cycle progression pathways, indicating a potent suppression of proliferative capacity. These transcriptomic data substantiate the MAESTRO-predicted mechanism: berberine acts as a synergistic driver by coupling the systemic collapse of mTORC1/AMPK signaling to a downstream transcriptional program of cell cycle arrest.

## Discussion

The development of effective combination therapies depends on deciphering the complex interplay between signaling pathway crosstalk and the core survival hubs that drive adaptive resistance. Identifying synergistic vulnerabilities within these non-linear networks requires an analytical scalability that exceeds the limits of manual curation. The MAESTRO framework addresses this challenge by leveraging the reasoning capabilities of LLMs, which have demonstrated significant problem-solving utility across diverse domains^61^, to transition from correlational synergy predictions toward a mechanism-driven discovery process. Central to this approach is the translation of LLM reasoning into quantifiable, uncertainty-aware rankings via an ensemble-consensus strategy. Consistent with previous studies, we demonstrate that model stochasticity is not merely noise, but a deconvolutable signal; high internal consensus across multiple reasoning paths indicates a semantic convergence on accurate biological mechanisms^62^. This "High-Confidence Zone" methodology aligns with current trends in high-stakes biomedical AI^63,64^, where generative consistency serves as a computational proxy for mechanistic accuracy. Ultimately, this paradigm enables the rapid screening of extensive bioactive libraries with high biological accuracy, utilizing the LLM’s reasoning power to overcome the complexities of therapeutic resistance. The framework’s ability to decipher signaling architecture proved essential for characterizing the molecular drivers of resistance.

Beyond broad pathway associations, the model demonstrated an "intra-pathway" resolution that differentiated between inhibitors of central survival hubs and those targeting peripheral downstream effectors. This prioritization of core signaling nodes aligns with established mechanistic principles of adaptive resistance, which suggest that targeting primary compensatory axes is required to mitigate the reciprocal feedback loops that sustain tumor cell viability^6^. Within this landscape, the platform identified the synergistic potential of modulating the PI3K/AKT and AMPK axes; while the reciprocal regulation between these pathways is mechanistically coupled^65^, their simultaneous pharmacological targeting remains a novel strategy in the context of therapeutic escape. This logic further facilitated the deconvolution of Berberine’s polypharmacology^66^, accurately predicting its ability to re-sensitize resistant cells by simultaneously attenuating PI3K/AKT/mTOR signaling and activating the AMPK-mediated metabolic stress axis.

Despite MAESTRO’s predictive utility, several constraints offer opportunities for systematic refinement. A primary limitation is the platform’s current reliance on text-based retrieval, which links inferential capacity to the breadth of existing literature and may reduce accuracy for novel chemical entities. Transitioning the framework toward de novo discovery will require the integration of molecular representations, such as SMILES strings or graph embeddings, directly into the model context^67^. Furthermore, the reliance on closed-source architectures introduces challenges regarding operational costs and longitudinal reproducibility. Future iterations utilizing high-parameter open-weight models could mitigate these issues, enabling domain-specific fine-tuning on specialized omics datasets to optimize internal weights for specific biological contexts while ensuring computational consistency and independence from proprietary version updates^68^.

In conclusion, the MAESTRO platform demonstrates that the statistical consistency of LLM outputs can be utilized to quantify the reliability of mechanistic inferences in drug discovery. By bridging the gap between generative AI and experimental validation, this work identifies the berberine-enzalutamide synergy as a potent strategy for overcoming complex adaptive resistance. Beyond the specific context of enzalutamide resistance, this framework presents a versatile strategy for therapeutic discovery. Its immediate utility lies in leveraging the rich literature surrounding standard-of-care agents, such as chemotherapeutics or PARP inhibitors, to uncover overlooked combinatorial opportunities. Conversely, for emerging inhibitors with sparse mechanistic data, the platform creates an opportunity to integrate functional priors: small-scale screenings or CRISPR-derived dependency maps can establish the initial mechanistic backbone, allowing MAESTRO to subsequently extrapolate and prioritize the larger combinatorial space. Ultimately, the integration of interpretable machine learning with rigorous validation established here provides a scalable, rational methodology for discovering multi-target therapies in oncology and beyond.

## Acknowledgements

C.L. is supported by financial support from National Defense Medical University, Taiwan. H. H. is supported by the United States Department of Defense (Award Number: HT9425-25–1-0290). T.S. is supported by the National Institutes of Health (NIH)/National Cancer Institute (NCI) (R37CA240822, R01CA287669 and R01CA274978), Worldwide Cancer Research, the United States Department of Defense (Award Number: HT9425-24–1-0396) and Neuroendocrine Tumor Research Foundation, Inc. (Award No. 20245682). A.S.G. is supported by UCLA Prostate Cancer Specialized Programs of Research Excellence (SPORE) NCI P50 CA092131, Department of Defense PCRP award (HT94252310379), the Mike Slive Foundation for Prostate Cancer Research, the UC Cancer Research Coordinating Committee (C23CR5598), and the 2024 Larry & Sherry Benaroya Prostate Cancer Foundation Challenge Award. N.Y.C.L. is supported by NSF (CBET-2244760, CMMI-2029454, DBI-2325121) and National Institutes of Health (NIH) NIGMS MIRA award (R35GM146735).

## Author contributions statement

C.H.L., A.S.G., and N.Y.C.L. conceived the projects; C.H.L., L.E., Y.H., C.J.H., and N.Y.C.L. developed the computational framework; C.H.L., K.H., L.K., A.B., J.D., Z.C., and H.H. conducted the experiments; C.H.L., K.H., L.K., A.B., J.D., Z.C., H.H., W.Y., A.L., R.D., T.S., A.S.G., and N.Y.C.L. analyzed the results. All authors reviewed the manuscript.

