## Supplemental Figure for "Mechanistic Language Modeling and Oxygenated 3D Screening Reveal Berberine and Enzalutamide Synergy in Resistant Prostate Cancer"

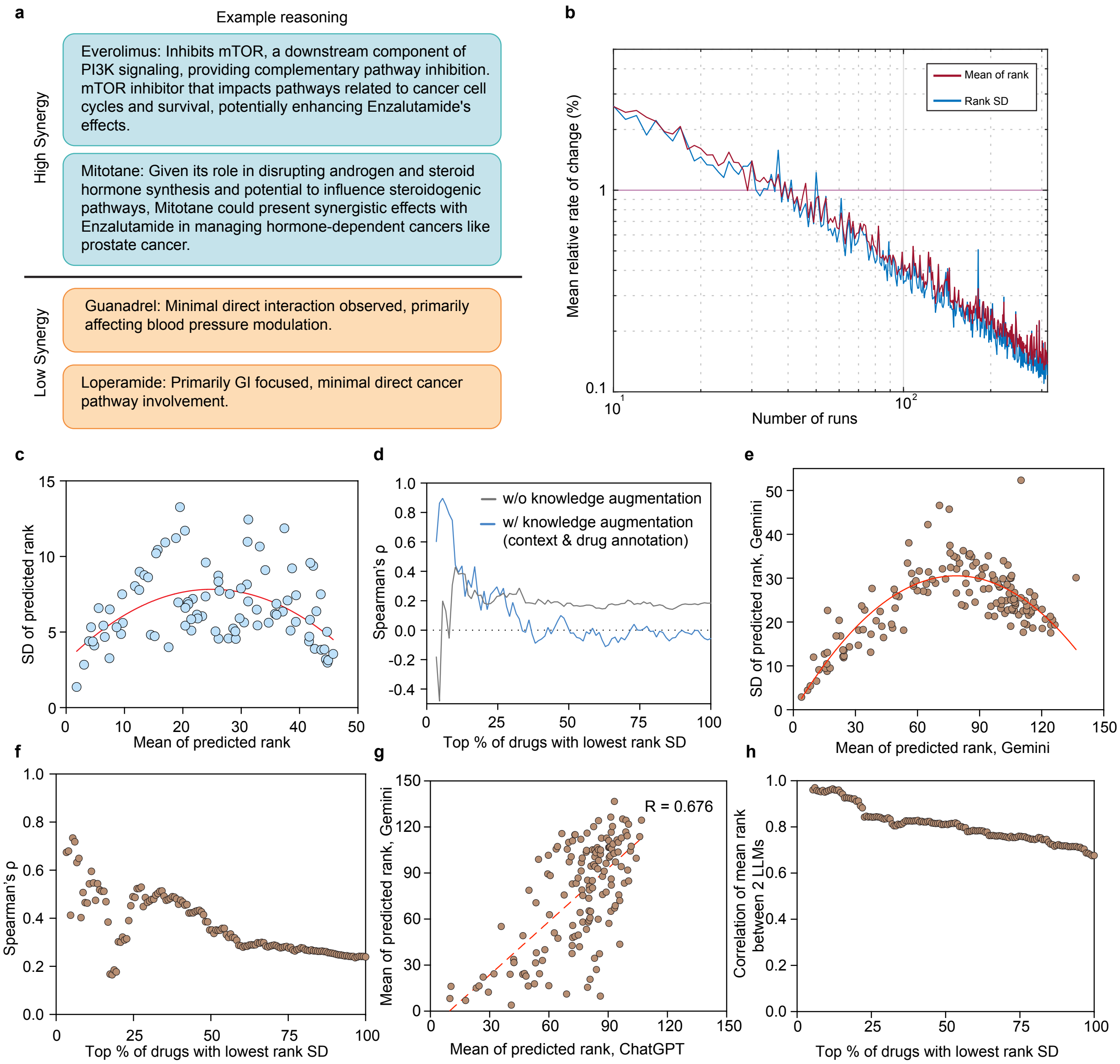


**Supplementary Figure 1. MAESTRO demonstrates generalization capabilities and cross-model consistency.** (**a**) Examples of MAESTRO-generated mechanistic reasoning distinguishing between high-synergy and low-synergy predictions. (**b**) Plot of mean relative rate of change (%) versus the number of runs per compound. (**c**) Scatter plot of MAESTRO-predicted rank standard deviation (SD) versus mean rank for 89 abiraterone-anchored drug pairs on the PC3 cell line using ChatGPT 4o. Red lines represent second-order polynomial (quadratic) fits. (**d**) Spearman's rank correlation coefficient (ρ) between predicted and experimental rankings as a function of the percentage of drugs included (ordered by increasing rank SD) for the dataset shown in (**c**). The plot presents an ablation analysis comparing model performance with and without knowledge augmentation. (**e**) Scatter plot of MAESTRO-predicted rank SD versus mean rank for 150 AR inhibitor-anchored drug pairs predicted using Google Gemini 2.5 Flash. Red lines represent second-order polynomial (quadratic) fits. (**f**) Spearman's rank correlation (ρ) between predicted and experimental rankings plotted against the percentage of drugs included (sorted by ascending rank SD) for the dataset in (**e**). (**g**) Scatter plot comparing mean ranks predicted by Gemini Flash (from panel (**e**)) versus ChatGPT (from **Fig. 1b**) for the identical set of 150 AR inhibitor-anchored drug pairs. The red dashed line represents the simple regression line. (**h**) Spearman's rank correlation (ρ) of mean rank between the two LLMs plotted against the percentage of drugs included (sorted by ascending rank SD), analyzing the stability of agreement shown in (**g**).

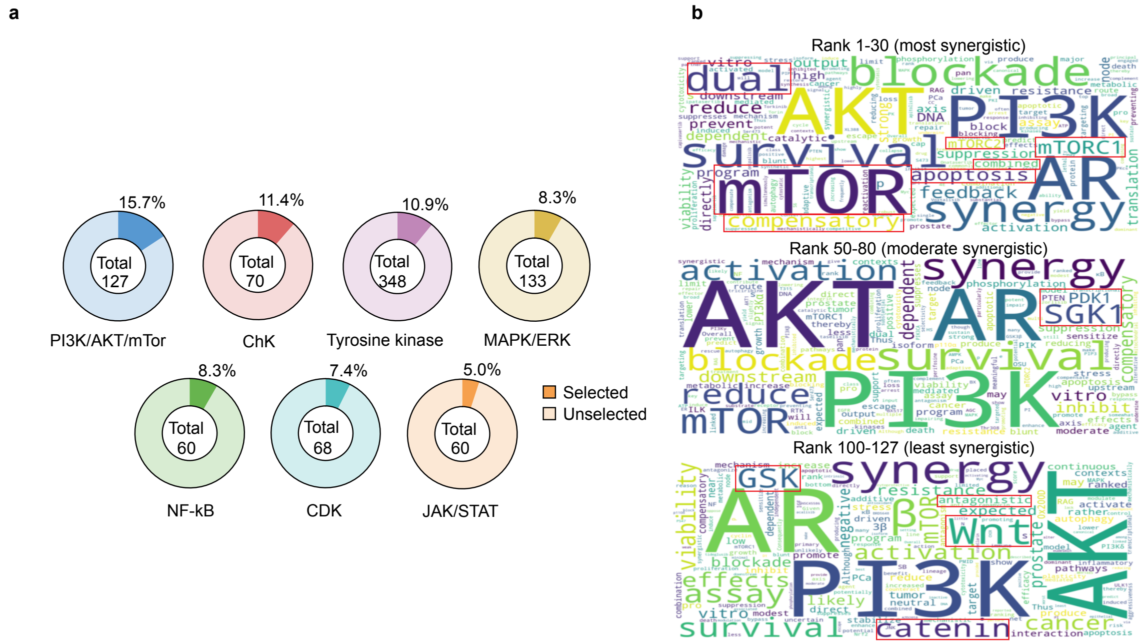


**Supplementary Figure 2. Distribution of mechanistic categories and text-mining analysis of MAESTRO predictions.** (**a**) Pie charts for each of the 7 mechanistic categories, displaying the total compound count (center) and the percentage of included compounds selected as one of the 95 prioritized candidates. (**b**) Word cloud analysis illustrating the frequency of terms in MAESTRO-generated mechanistic justification (100 runs per inhibitor) for 127 PI3K/AKT/mTOR inhibitors, stratified into three groups based on predicted mean rank. Text size is proportional to the frequency of term occurrence.

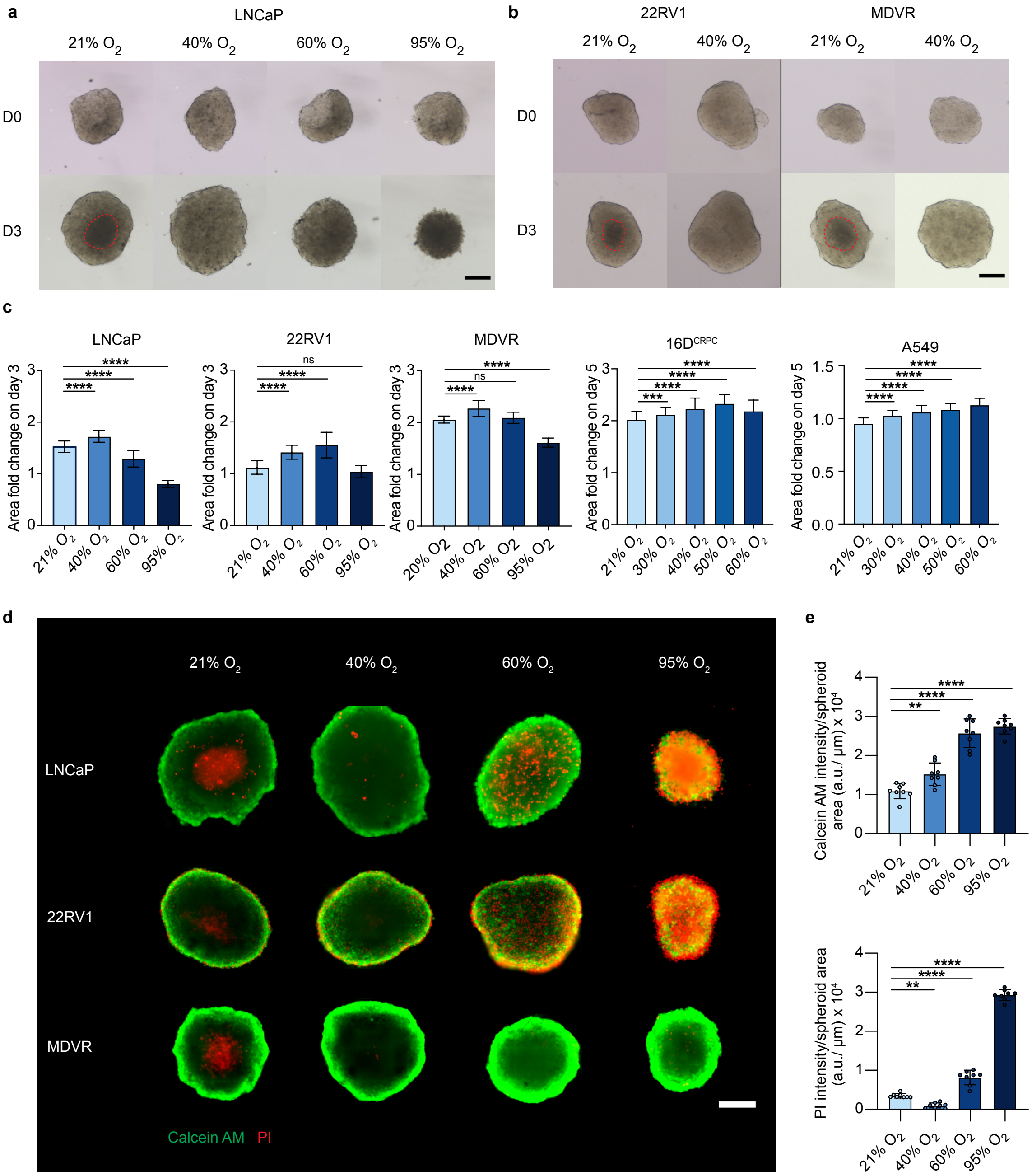


**Supplementary Figure 3. Spheroids exhibit robust growth kinetics and cellular viability under optimal oxygen supplementation.** (**a-b**) Representative bright-field images of (**a**) LNCaP, (**b**, left) 22RV1 , and (**b**, right) MDVR spheroids cultured under indicated oxygen levels 0 (top) and 3 (bottom) days after spheroid formation. (**c**) Bar charts showing spheroid area fold change at day 3 or 5 under indicated oxygen levels in LNCaP (n = 82-106), 22RV1 (n = 35-49), MDVR (n = 16-26), $16D^{CRPC}$ (n = 75-125), and A549 (n = 37-48) spheroids. (**d**) Representative Live/Dead staining images of LNCaP, 22RV1, and MDVR spheroids cultured under indicated oxygen levels. Green signal: Calcein AM (live cells); Red signal: Propidium Iodide (PI, dead cells). (**e**) Quantification of integrated Calcein AM (top) and PI (bottom) signal intensity normalized to spheroid area in LNCaP spheroids across different oxygen levels (n = 8). Scale bar, 200 µm. Data are presented as mean ± SD. Statistical significance was determined by one-way ANOVA followed by Dunnett's multiple comparisons test; * $P < 0.05$, ** $P < 0.01$, *** $P < 0.001$, **** $P < 0.0001$, ns: not significant.

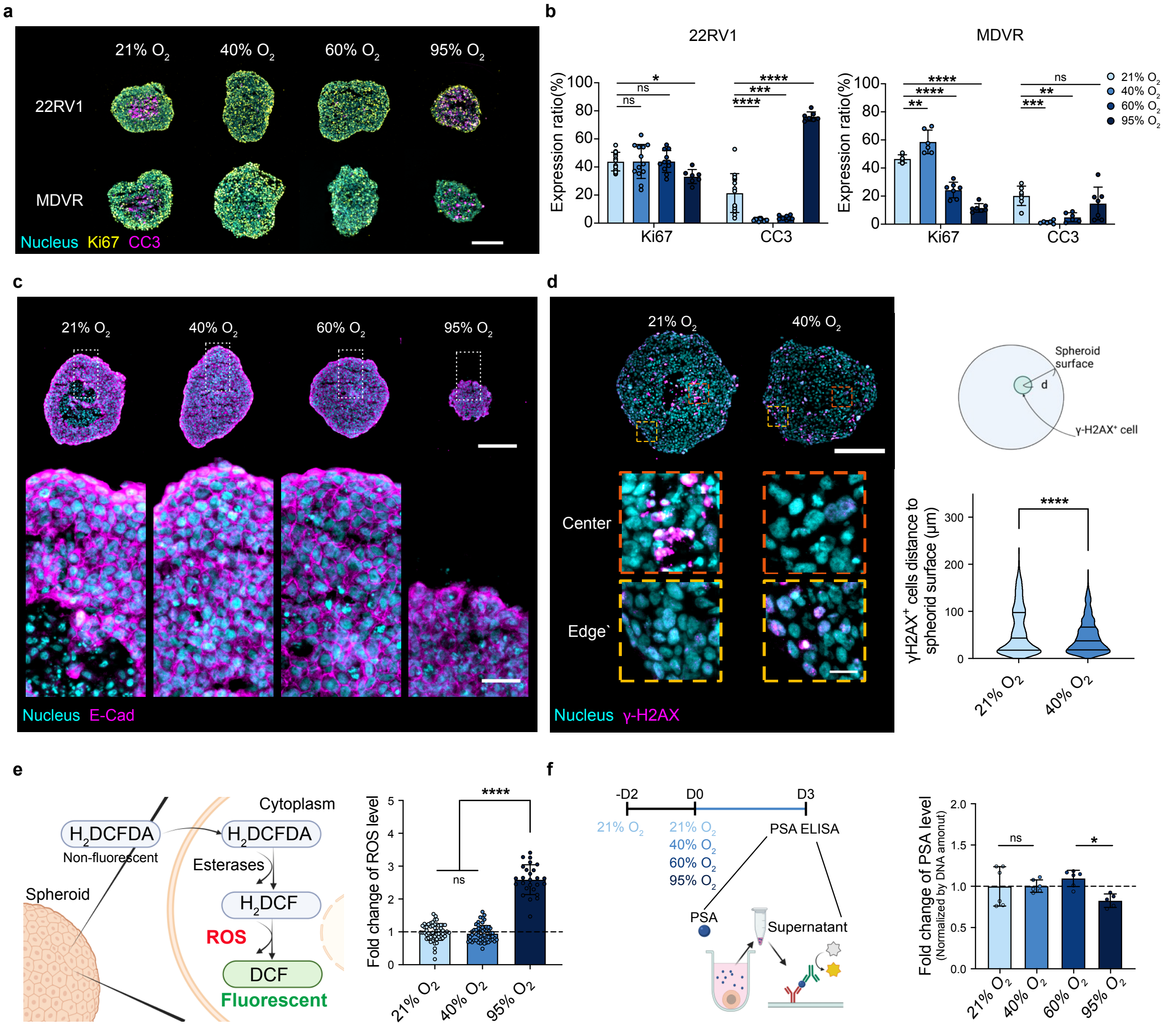


**Supplementary Figure 4. Spheroids maintain structural integrity and physiological function under optimal oxygen supplementation.** (**a-b**) Representative immunofluorescence images of (**a**) Ki67 (proliferation marker; yellow) and cleaved caspase-3 (CC3; apoptosis marker; magma) in 22RV1 (n =7-15) and MDVR (n = 6-7) spheroids cultured under indicated oxygen levels and (**b**) corresponding quantification. Scale bar, 200 µm. (**c**) Representative E-cadherin (E-cad) staining of LNCaP spheroids illustrating cell junction integrity. Bottom panels display magnified views of the regions outlined in white, spanning from the spheroid edge to the spheroid core. Scale bars, 200 µm (top) and 40 µm (bottom). (**d**) Representative immunofluorescence staining of γ-H2AX (DNA double-strand break marker) in LNCaP spheroids cultured under 21% and 40% $O_2$ (left). Bottom panels show magnified views of the spheroid core and edge regions. Violin plot (right, bottom) quantifies the shortest distance from each γ-H2AX+ cell to the spheroid surface (n = 979 cells for 21% $O_2$; n = 675 cells for 40% $O_2$). Scale bars, 200 µm (top) and 20 µm (bottom). (**e**) Schematic illustrations (left) and corresponding quantification (right) of intracellular ROS levels using $H_2$DCFDA (n = 56 for 21% and 40% $O_2$; n = 26 for 95% $O_2$). (**f**) Secreted PSA levels quantified via ELISA in LNCaP spheroids on day 3 of spheroid culture (left). Bar charts (right) present values normalized to total DNA content and relative to the 21% $O_2$ control (n = 6). Data are presented as mean ± SD. Statistical significance was determined by one-way ANOVA followed by Dunnett's multiple comparisons test (**b, e, f**) or two-tailed unpaired Student's t-test (**d**); * $P < 0.05$, ** $P < 0.01$, *** $P < 0.001$, **** $P < 0.0001$, ns: not significant.

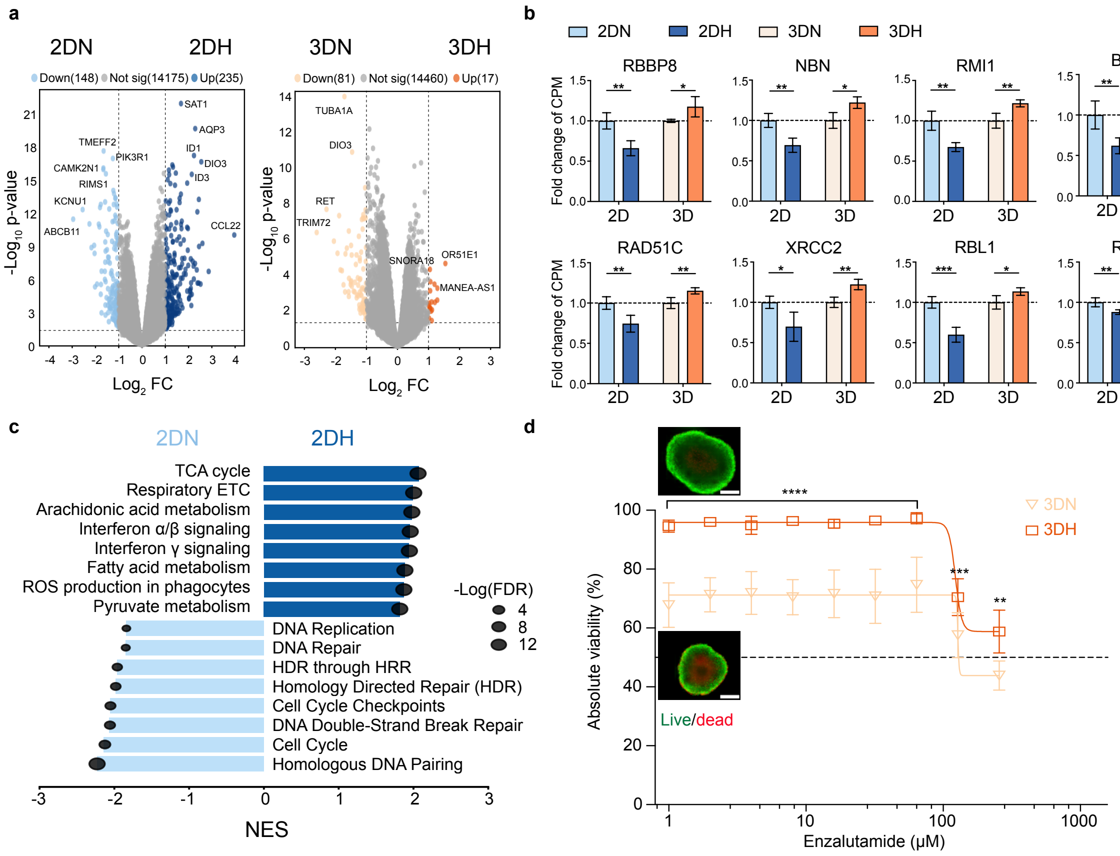


**Supplementary Figure 5. Oxygen supplementation modulates transcriptomic profiles and drug sensitivity in prostate cancer spheroids.** (**a**) Volcano plots displaying differential gene expression in LNCaP cells comparing 2D normoxia (2DN; 21% $O_2$) vs. 2D hyperoxia (2DH; 40% $O_2$) and 3D normoxia (3DN) vs. 3D hyperoxia (3DH). Dotted lines indicate significance thresholds ($|\log_2FC| \geq 1$, $P < 0.05$; n = 4). (**b**) Bar charts showing representative genes related to DNA repair and cell cycle pathways. Oxygen supplementation led to downregulation of these genes in 2D cultures, whereas this effect was reversed in 3D spheroids (n = 4). (**c**) Gene Set Enrichment Analysis (GSEA) of upregulated and downregulated pathways in the 2DN versus 2DH comparison (n = 4). (**d**) Enzalutamide dose-response curves plotting absolute viability (from Live/Dead staining) for 22RV1 spheroids in 3DN and 3DH groups. Insets display representative images of untreated controls at Day 3 (n = 6). Scale bar, 200 μm. Data are presented as mean ± SD. Statistical significance was determined by two-tailed unpaired Student's t-test; * $P < 0.05$, ** $P < 0.01$, *** $P < 0.001$, **** $P < 0.0001$, ns: not significant.

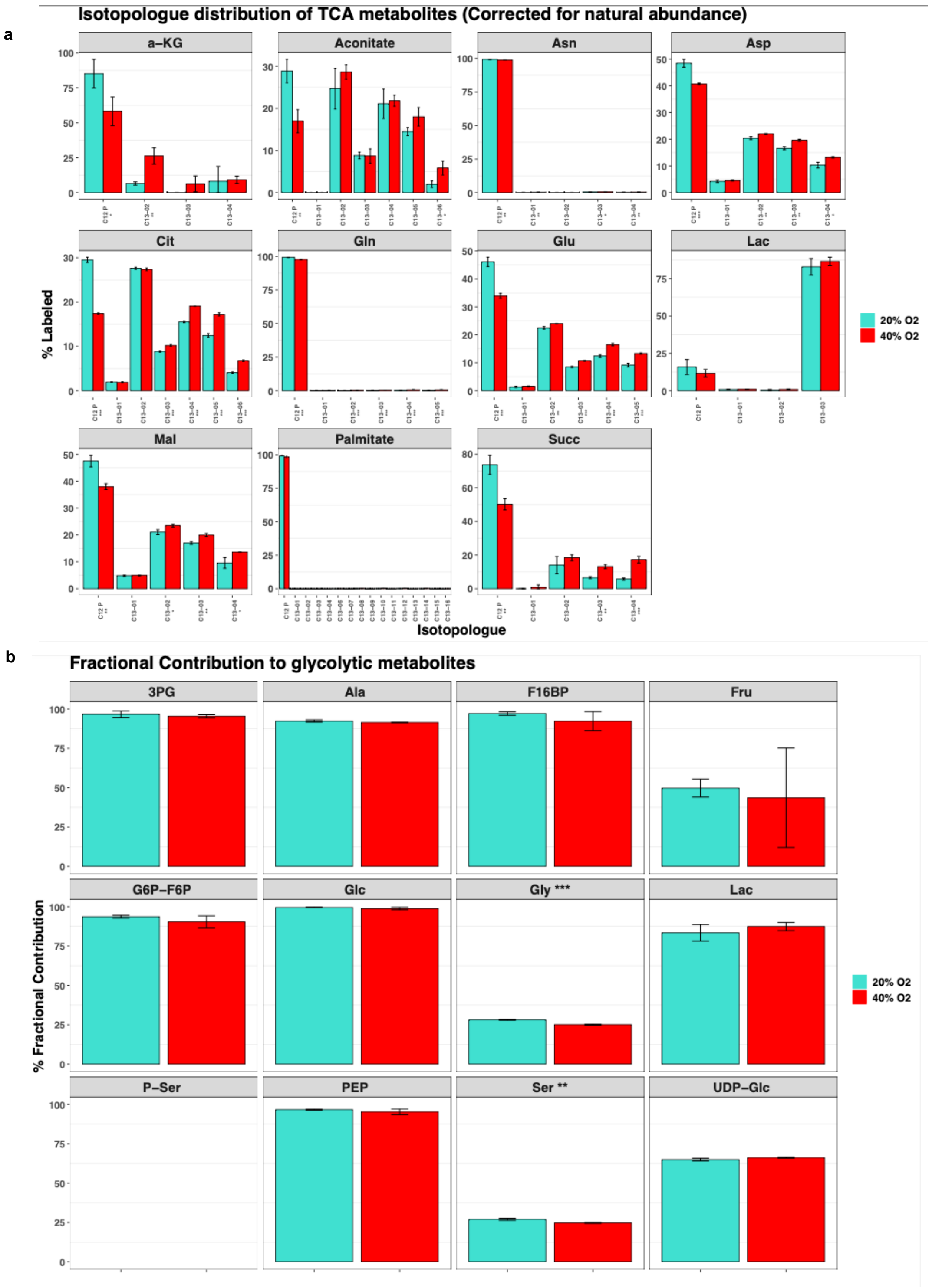


**Supplementary Figure 6. Metabolomic profiling reveals that oxygen supplementation promotes glucose entry into the TCA cycle without altering glycolysis.** (**a**) Isotopologue distribution of TCA cycle metabolites in LNCaP spheroids cultured under indicated oxygen levels. (**b**) Fractional contribution to glycolytic metabolites in LNCaP spheroids cultured under indicated oxygen levels. Spheroids were labeled with D-[U-$^{13}C_6$] glucose for 24 h prior to analysis. Data are presented as mean ± SD (n = 3). Statistical significance was determined by two-tailed unpaired Student's t-test; * P<0.05, ** P<0.01, *** P<0.001, **** P<0.0001, ns: not significant.

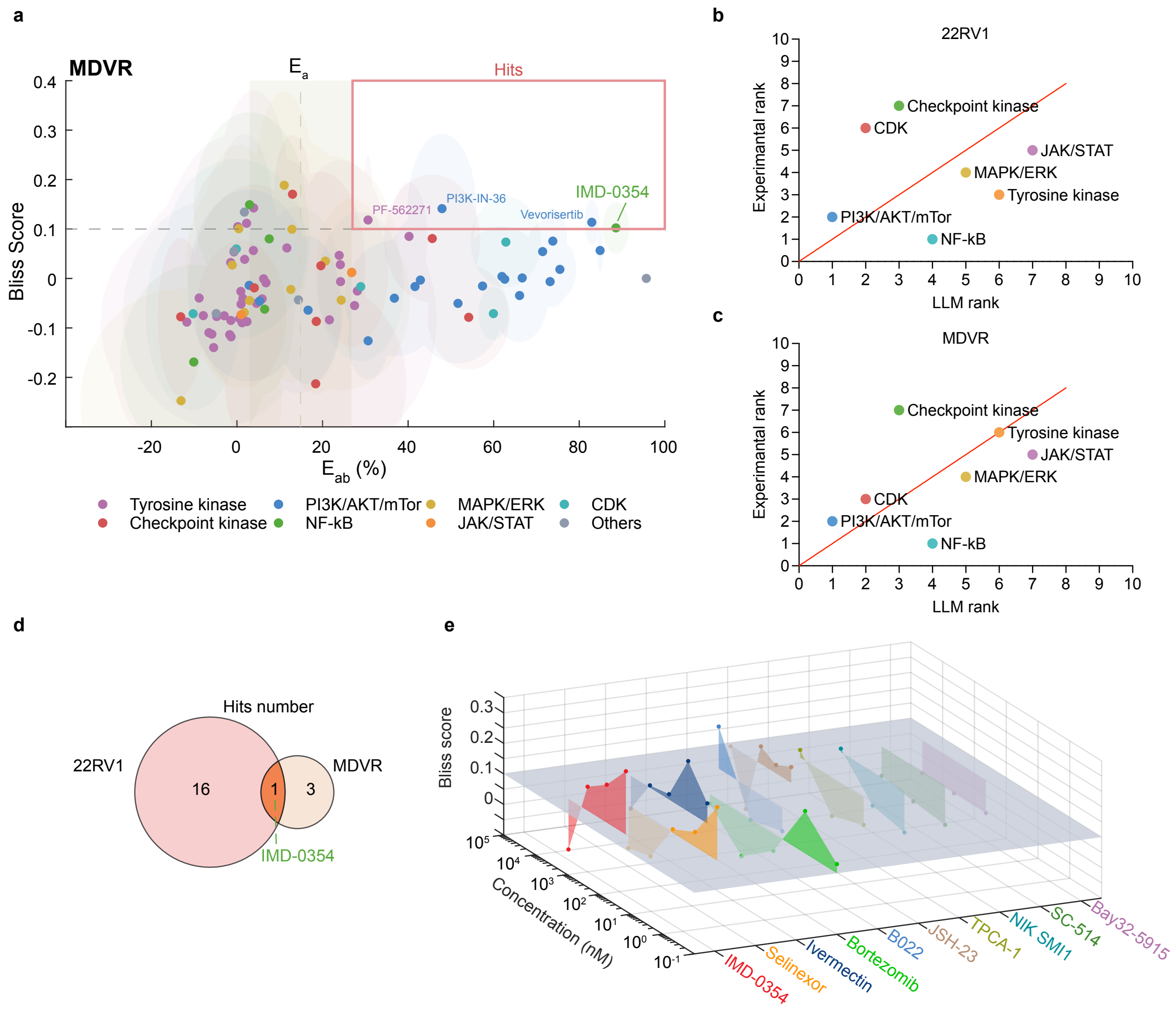


**Supplementary Figure 7. In vitro screening validates the pathway-specific predictive accuracy of MAESTRO-prioritized PKIs.** (**a**) Scatter plot of Bliss score versus $E_{ab}$ for 95 PKIs tested in oxygenated MDVR spheroids. Hits (red boxed region) are defined as compounds with $E_{ab}$ greater than enzalutamide monotherapy ($E_a$) and a Bliss score > 0.1. Error ellipses denote the SD (n = 4) for $E_{ab}$ (x-axis) and Bliss scores (y-axis). (**b-c**) Scatter plots displaying the relationship between experimental rank and category-averaged MAESTRO rank (LLM rank) across 7 mechanistic categories in (**b**) 22RV1 and (**c**) MDVR spheroids cultured under 40% $O_2$. (**d**) Venn diagram displaying hit numbers identified in oxygenated 22RV1 and MDVR spheroids and their shared hit, IMD-0354. (**e**) Three-dimensional plot displaying Bliss scores for 10 NF-κB inhibitors across multiple concentrations.

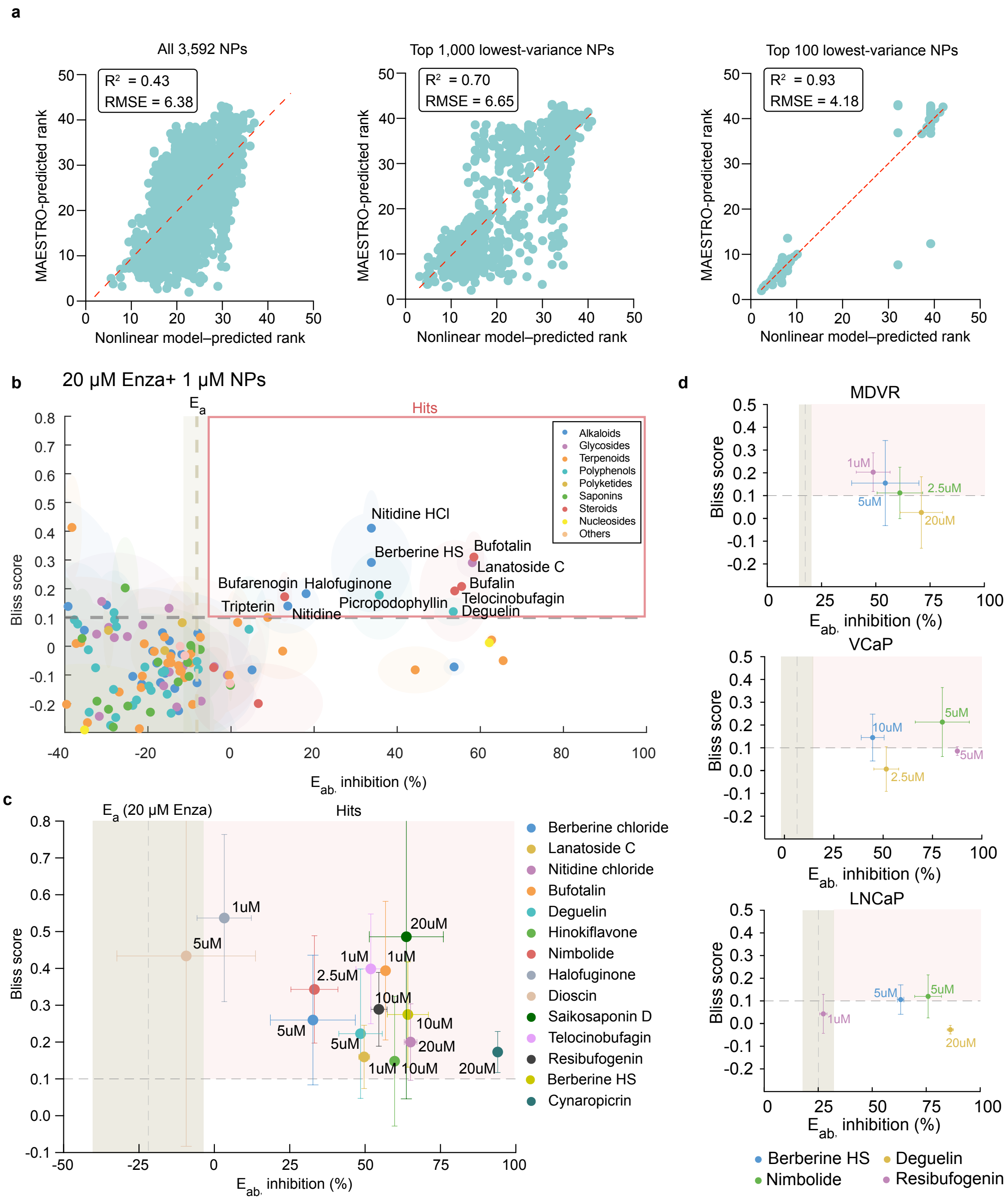


**Supplementary Figure 8. Multi-stage screening identifies MAESTRO-predicted natural products as potent synergistic agents.** (**a**) Correlation plots illustrating the MAESTRO-predicted mean rank versus the nonlinear GBT model-predicted rank. Panels represent the full dataset (left), top 1,000 candidates (middle), and top 100 lowest-variance (high confidence) candidates (right). Red dashed lines indicate linear regression fits. (**b**) Screening of 150 prioritized candidates in MDVR spheroids plotting combination efficacy ($E_{ab}$) against Bliss synergy scores. Hits (red boxed region) are defined as compounds exhibiting $E_{ab} > E_a$ and a Bliss score > 0.1. Error ellipses and shaded band represent SD (n = 4). (**c**) Validation of 14 top-tier candidates showing Bliss score versus $E_{ab}$. Labels indicate the concentration yielding the highest Bliss score for each compound (n = 4). (**d**) Synergy validation of berberine hydrogen sulphate, deguelin, nimbolide, and resibufogenin across MDVR (top), VCaP (middle), and LNCaP (bottom) spheroids (n = 8). In (**b–d**), the vertical dashed lines and shaded areas represent the mean ± SD of enzalutamide monotherapy efficacy ($E_a$). Data are presented as mean ± SD.
